# A defined clathrin-mediated trafficking pathway regulates sFLT1/VEGFR1 secretion from endothelial cells

**DOI:** 10.1101/2023.01.27.525517

**Authors:** Karina Kinghorn, Amy Gill, Allison Marvin, Renee Li, Kaitlyn Quigley, Ferdinand le Noble, Feilim Mac Gabhann, Victoria L Bautch

**Affiliations:** Curriculum in Cell Biology and Physiology, University of North Carolina, Chapel Hill NC USA; Department of Biology, University of North Carolina, Chapel Hill NC USA; McAllister Heart Institute, University of North Carolina, Chapel Hill NC USA; UNC Lineberger Comprehensive Cancer Center, University of North Carolina, Chapel Hill NC USA; Department of Cell and Developmental Biology, Institute of Zoology, Karlsruhe Institute of Technology, Karlsruhe, Germany; Institute for Computational Medicine and Department of Biomedical Engineering, Johns Hopkins University, Baltimore MD, USA

**Author notes:** Corresponding author: Victoria L. Bautch, PhD, Department of Biology, CB No. 3280, The University of North Carolina at Chapel Hill Chapel Hill, NC 27599 USA.

**Keywords:** sFLT1, secretion, Golgi, AP1, clathrin, vWF

## Abstract

FLT1/VEGFR1 negatively regulates VEGF-A signaling and is required for proper vessel morphogenesis during vascular development and vessel homeostasis. Although a soluble isoform, sFLT1, is often mis-regulated in disease and aging, how sFLT1 is trafficked and secreted from endothelial cells is not well understood. Here we define requirements for constitutive sFLT1 trafficking and secretion in endothelial cells from the Golgi to the plasma membrane, and we show that sFLT1 secretion requires clathrin at or near the Golgi. Perturbations that affect sFLT1 trafficking blunted endothelial cell secretion and promoted intracellular mis-localization in cells and zebrafish embryos. siRNA-mediated depletion of specific trafficking components revealed requirements for RAB27A, VAMP3, and STX3 for post-Golgi vesicle trafficking and sFLT1 secretion, while STX6, ARF1, and AP1 were required at the Golgi. Depletion of STX6 altered vessel sprouting in a 3D angiogenesis model, indicating that endothelial cell sFLT1 secretion is important for proper vessel sprouting. Thus, specific trafficking components provide a secretory path from the Golgi to the plasma membrane for sFLT1 in endothelial cells that utilizes a specialized clathrin-dependent intermediate, suggesting novel therapeutic targets.

## INTRODUCTION

Blood vessels form early during vertebrate embryonic development to deliver oxygen and nutrients to developing tissues and organs, and in adults blood vessels also regulate organ function and homeostasis [3–5]. Endothelial progenitor cells initially coalesce to create vessels that differentiate through a process called vasculogenesis, and new vessels then arise from existing vessels primarily via sprouting angiogenesis to expand the vascular network [6–8]. Numerous signaling pathways regulate angiogenic sprouting and cross-talk with each other, and signaling amplitude is often controlled by endothelial cell-intrinsic negative regulators of the pathways. Thus, endothelial cells are crucial players in angiogenic pathway regulation during blood vessel formation and function.

Among the signals important to sprouting angiogenesis, vascular endothelial growth factor-A (VEGF-A)-mediated signaling stands out, as it is required in almost all tissues for proper blood vessel formation [9–11]. The VEGF-A ligand is alternatively spliced to produce several isoforms that differentially interact with the extracellular matrix (ECM) and thus provide spatial cues and survival signals to endothelial cells in emerging vessels [12]. Endothelial cells often respond to VEGF-A signals by adopting a migratory tip cell phenotype or a more proliferative stalk cell phenotype to extend the sprout [13,14]. Among endothelial cell VEGF-A receptors, VEGFR2 (FLK1) and VEGFR1 (FLT1) provide the primary pro- and anti-angiogenic signals, respectively. Tip cells express higher levels of VEGFR2 that promote migration, while stalk cells express higher levels of FLT1 [15,7]. FLT1 is alternatively spliced to produce a full-length transmembrane isoform (mFLT1) and a secreted soluble isoform (sFLT1) [16,17]. Both FLT1 isoforms bind VEGF with a 10-fold higher affinity than VEGFR2 and can act as decoys to sequester excess VEGF-A ligand [18,19]. FLT1, VEGFR2, and VEGF-A are all required for blood vessel formation, as genetic loss leads to severe vascular defects and early embryonic lethality in mice [20–23]. However, FLT1 signaling via its cytoplasmic tyrosine kinase domain is not required for vascular development or viability, as mice lacking the cytoplasmic portion of FLT1 survive without defects in vascular development, and mice expressing only sFLT1 have significant viability [24,25]. sFLT1 secretion is upregulated in response to hypoxia and is associated with endothelial cell dysfunction during aging, chronic kidney disease, and COVID-19 infection [26–29]. Thus, negative regulation of VEGF-A signaling through FLT1 is required for proper vascular development, and dysregulation is associated with vascular pathology and aging.

sFLT1 is secreted from endothelial cells and binds the ECM via poorly-defined heparin-binding sites, thus acting as a critical molecular rheostat to modulate VEGF-A availability extracellularly [30]. In mouse embryonic stem cell-derived *Flt1−/−* mutant vessels, genetic rescue with either endothelial cell-expressed *mFlt1* or *sFlt1* restored proper VEGFR2 signaling and endothelial cell proliferation, but only *sFlt1* effectively rescued branching morphogenesis [31]. Additionally, *sFlt1* expression from the lateral base area of new sprouts restored the ability of emerging sprouts to extend away from the parent vessel that was lost with *Flt1* deletion [32,33]. These findings support a local sprout guidance model, positing that stalk cell secretion of sFLT1 neutralizes VEGF-A next to the sprout base, thus establishing a forward guide for tip cells as they migrate away from the parent vessel. In zebrafish, genetic loss of both *mflt1* and *sflt1* leads to ectopic sprouting of intersegmental vessels, while expression of only *sflt1* was sufficient for proper vessel formation [34,35]. Together, these data suggest that secreted sFLT1 interactions in the ECM are critical for regulating blood vessel development and patterning. Despite this required function for sFLT1, the mechanisms regulating sFLT1 trafficking and secretion from endothelial cells remain largely unknown.

Secreted proteins are most often trafficked through the endoplasmic reticulum (ER) to the cis- and then the trans-Golgi, where they are loaded into vesicles for transport to the plasma membrane and secretion [36]. Trafficking and secretion are largely regulated by three classes of proteins: ARF (ADP-ribosylation factor) GTPases, RAB GTPases, and SNAREs (Soluble NSF Attachment Protein Receptors) [37–41]. Within this general framework, multiple molecular components contribute to three major pathways involved in secretion: unregulated secretion via vesicles, trafficking via an intermediate sorting compartment, and stimulus-regulated release from large storage granules that fuse with the plasma membrane [42]. In endothelial cells, a well-characterized secretory pathway leads to von Willebrand Factor (vWF) release from Weibel-Palade bodies upon histamine stimulation to participate in the clotting cascade [43]. These large storage granules are held at the plasma membrane until stimulus-promoted fusion and extracellular release of contents [44–47]. Absent stimulation, vWF is constitutively secreted through Golgi-derived vesicles or via spontaneous release of Weibel-Palade bodies [48,49] using poorly defined pathways that are distinct from stimulated vWF release. Although numerous other proteins are associated with Weibel-Palade bodies, sFLT1 has not been detected in Weibel-Palade bodies [50], suggesting a unique trafficking pathway.

Although sFLT1 is constitutively secreted from endothelial cells, the trafficking pathway from the ER to the cell surface for sFLT1 has not been well-defined. sFLT1 is detected in human plasma and conditioned media of endothelial cells purified from different organs absent stimulation [51–53], and secretion is stimulated by hypoxia [54–56]. Stable overexpression of tagged sFLT1 in endothelial cells revealed a requirement for ARF1, RAB11, and ARF6 in trafficking of the labeled protein [2], but no studies have systematically defined how sFLT1 is trafficked through endothelial cells to the plasma membrane for secretion.

Here, we define a constitutive trafficking pathway for sFLT1 secretion from endothelial cells. Surprisingly, constitutive sFLT1 secretion utilizes trafficking components that are typically associated with stimulated secretion of Weibel-Palade bodies, such as AP1-dependent clathrin assembly and RAB27a [50]. Moreover, the SNARE STX6, which is typically involved with Golgi-to-endosomal trafficking, is required for sFLT1 secretion. Perturbations that affect secretion also mis-localize sFLT1 in endothelial cells *in vitro* and *in vivo,* and affect 3D vascular sprouting. Thus, endothelial cell sFLT1 is trafficked by several regulators prior to its secretion, and this pathway is novel in its entirety, suggesting new therapeutic targets for treatment of endothelial cell dysfunction.

## RESULTS

### sFLT1 is constitutively secreted from endothelial cells

Although it is well-established that soluble FLT1 (sFLT1) is secreted from endothelial cells [16,2], the pathway utilized by sFLT1 for trafficking and secretion is poorly described. To define a pathway from synthesis to secretion in endothelial cells, we first confirmed that sFLT1 is robustly secreted from primary human umbilical vein endothelial cells (HUVEC) absent stimulation. Concentrated conditioned media and lysates of HUVEC but not normal human lung fibroblast (NHLF) controls contained a 90 kDa protein that reacted with a FLT1 antibody **(Fig. 1A).** Selective manipulation of the two FLT1 isoforms, mFLT1 and sFLT1, via siRNA depletion revealed that siRNA targeting both FLT1 isoforms (si*FLT1*) reduced both 180 kDa mFLT1 and 90 kDa sFLT1 protein levels, while siRNA that selectively targeted sFLT1 (si*sFLT1*) [57] reduced the 90 kDa sFLT1 protein levels in media and lysates but not the 180 kDa mFLT1 protein in lysates (**Fig. 1B)**. Thus, sFLT1 is constitutively secreted by endothelial cells and accumulates to detectable intracellular levels in HUVEC.

**Fig. 1.**
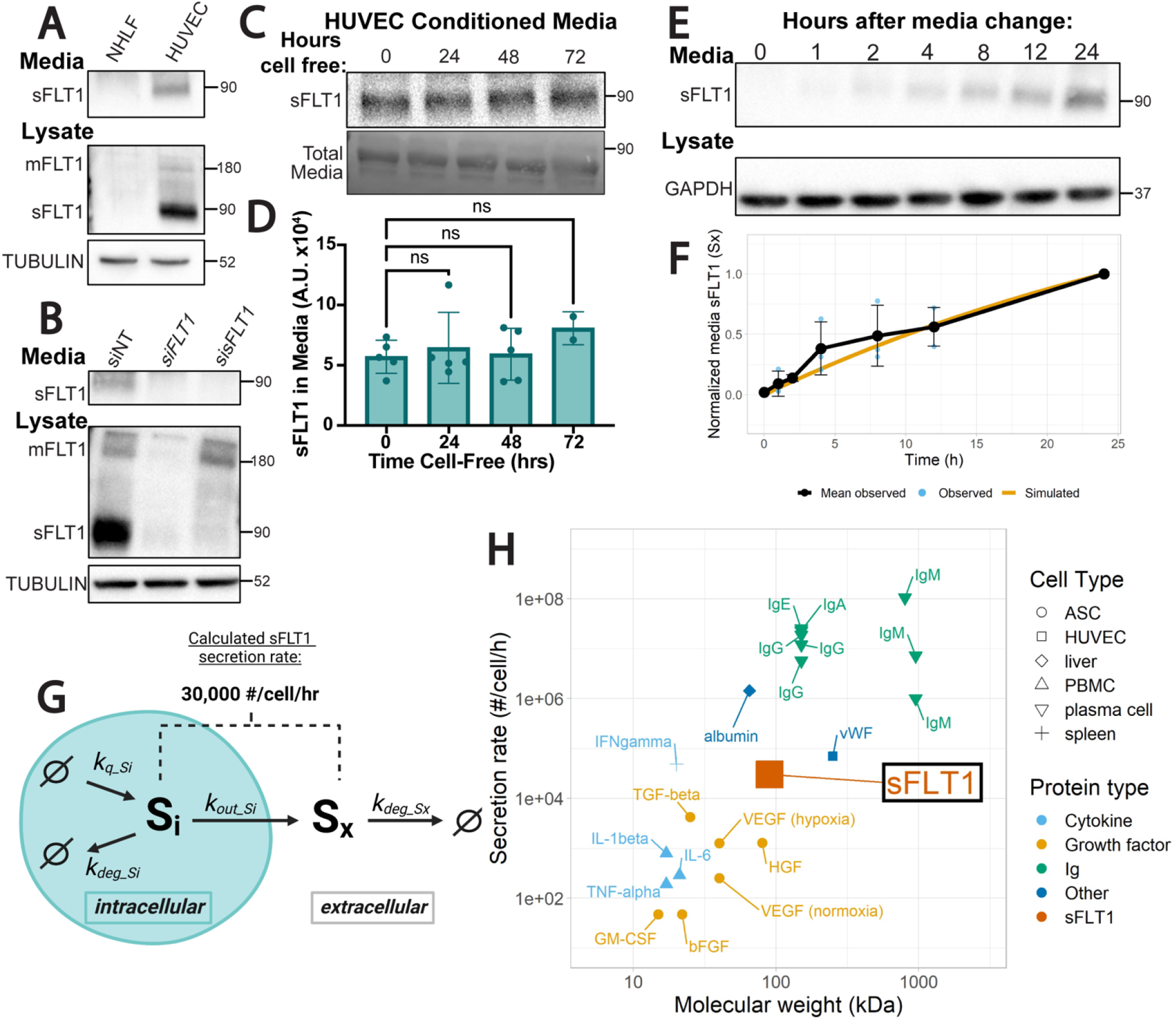
sFLT1 is constitutively secreted from endothelial cells. (A) Immunoblot of FLT1 in concentrated conditioned media or cell lysates of normal human lung fibroblasts (NHLF) or human umbilical vein endothelial cells (HUVEC). sFLT1 detected at 90 kDa and mFLT1 detected at 180 kDa. Tubulin loading control. Representative of 3 replicates. (B) Immunoblot of FLT1 in concentrated media or lysates of HUVEC transfected with siRNAs targeting mFLT1 and sFLT1 (si*FLT1*), sFLT1 (si*sFLT1*), or non-targeting control (siNT). Tubulin loading control. Representative of 3 replicates. (C) Top: representative immunoblot of sFLT1 in concentrated conditioned media from HUVEC incubated cell-free for indicated times. Bottom: total media protein loading control. (D) Quantification of FLT1 band intensity from 1C. Shown are means +/− SD of n=2-5 replicates/condition. (E) Representative immunoblot of sFLT1 in HUVEC concentrated conditioned media for the indicated time points following media change. GAPDH loading control. (F) Graph of sFLT1 media/time. Black line: densitometric analysis of sFLT1 from 1E, with media sFLT1 normalized to intensity at 24h (n=3 replicates). Blue dots and error bars: Means +/− SD of n=3 replicates/condition. Orange line: simulation of secreted sFLT1 from literature-based computational model [1,2]. **(G)** Diagram of processes included in the mechanistic computational model of sFLT1 secretion, with the calculated secretion rate noted. Schematic created with BioRender.com. **(H)** Comparison of sFLT1 secretion rate predicted by computational model vs. molecular weight to that of other secreted proteins (See Supplementary Table S5). Abbreviations: ASC = adipose stromal cell, HUVEC = human umbilical vein endothelial cell, Ig = immunoglobulin, PBMC = peripheral blood mononuclear cell.

To determine the basal rate of sFLT1 secretion from endothelial cells, we analyzed sFLT1 protein levels in media and lysate collected from HUVEC over time. We first assessed the stability of extracellular sFLT1 and saw no significant decrease in levels up to 72h in media that was not exposed to cells **(Fig. 1C-D)**, allowing us to equate media protein levels with the amount of secreted protein over time **(Fig. 1E)**. We next quantified the levels over time after a media change and compared these data to predictions from a newly generated mechanistic computational model of sFLT1 secretion (**Fig. 1F-G**; **Methods; Supplementary Fig. 1A**). For the computational model, rate parameters were derived from published time course data of sFLT1 secretion (**Supplementary Tables S1-S4**). The model parameters are well-constrained by the previous data, with low uncertainty in the optimal values (**Supplementary Fig. 1B-D**) and also simulate the data in this study well (**Fig 1F**). As a result, we estimate that sFLT1 is constitutively secreted at a constant rate of approximately 30,000 molecules/cell/hr (**Fig. 1G**). This value is comparable to the constitutive secretion rate of vWF in HUVEC [58], faster than the published secretion rate of VEGF-A and other cytokines/growth factors from adipose stromal cells and peripheral blood mononuclear cells [59,60], and slower than published rates of immunoglobulins from plasma cells or albumin from liver cells [61–66] (**Fig. 1H**, **Supplementary Table S5**). These data are consistent with sFLT1 being produced and secreted at physiologically relevant levels by HUVEC, and confidence in the secretion estimate is enhanced by consistency across multiple independent experiments by different groups. The model also gives insight into relationships between mechanistic processes affecting sFLT1; for example, the modeling suggests that an intracellular sFLT1 molecule has a roughly equal probability of being secreted or intracellularly degraded (**Supplementary Table S4**).

### Intracellular sFLT1 is Golgi-localized prior to secretion

To begin defining how sFLT1 is trafficked in endothelial cells, we determined sFLT1 subcellular localization. C-terminal HA-tagged human sFLT1 (sFLT1-HA) was introduced to non-endothelial cells, and immunoblot analysis confirmed that both cell-associated and secreted sFLT1-HA were detected by HA and FLT1 antibodies (**Supp. Fig. 2A**). sFLT1-HA was also detected in conditioned media and lysate from transfected HUVEC with both HA and FLT1 antibodies (**Fig. 2A**). Immunofluorescence imaging of sFLT1-HA intracellular localization showed that in both non-endothelial cells and HUVEC, sFLT1-HA colocalized with the trans-Golgi marker, γ-Adaptin, consistent with previous reports [2,67] **(Fig. 2B, B’**, **Supp. Fig. 2B, B’)**. Staining for endogenous FLT1 revealed a similar perinuclear localization in HUVEC that colocalized with γ-Adaptin as shown by the line scan profile **(Fig. 2C, C’)**, and FLT1 antibody reactivity was specific to HUVEC and not NHLF controls **(Supp. Fig. 2C)**. We next examined the localization of sFLT1 by subcellular fractionation via density gradient ultracentrifugation of HUVEC lysates, and confirmed that most sFLT1 protein is found in fractions that also express the trans-Golgi marker, STX6, with lesser amounts seen in fractions that express the ER marker calnexin, and in vesicle/ plasma membrane fractions **(Fig. 2D)**. Therefore, prior to secretion from endothelial cells, sFLT1 is Golgi-localized, and minor depots of sFLT1 protein suggest it moves from the ER to the Golgi, and from the Golgi to cytoplasmic vesicles prior to secretion.

**Fig. 2.**
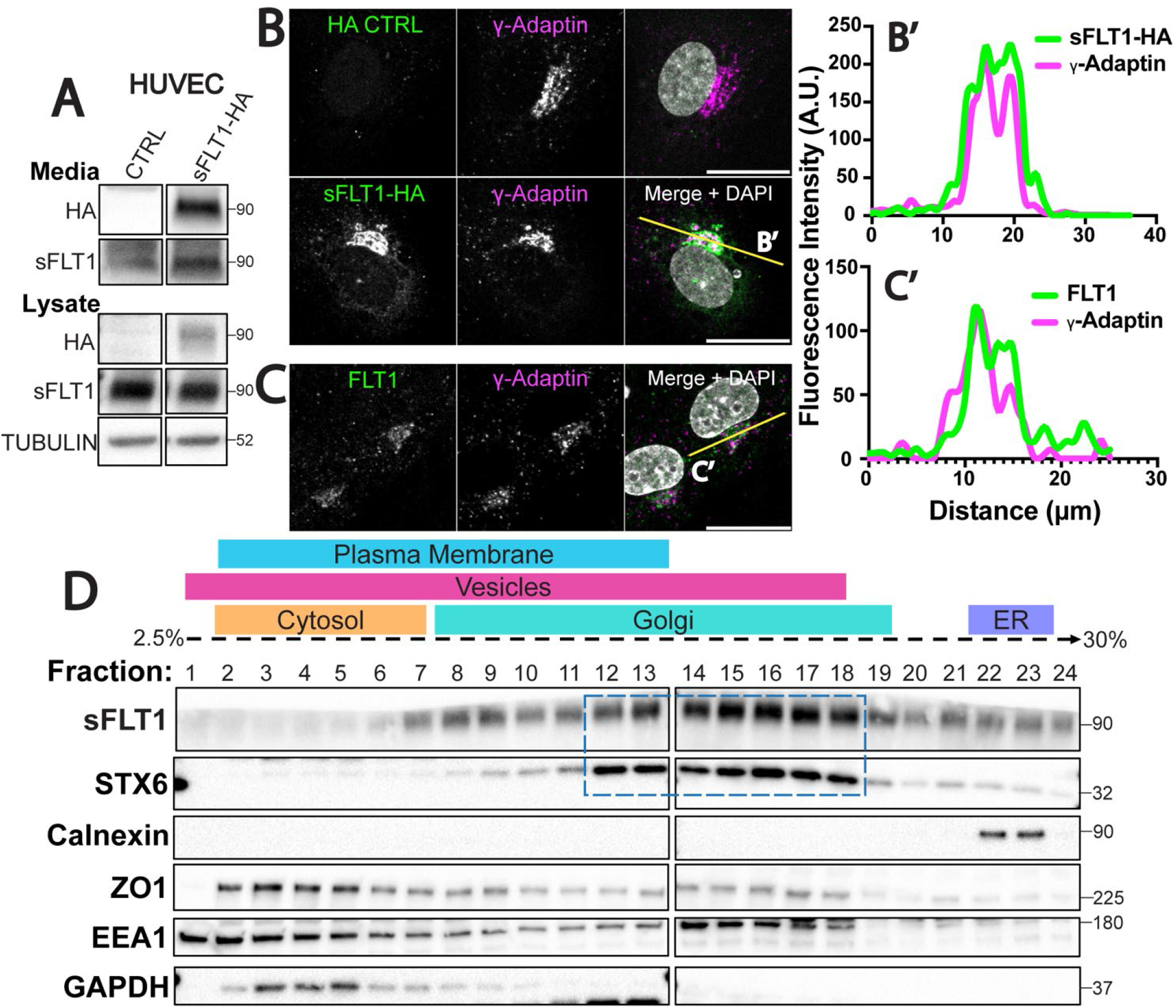
Intracellular sFLT1 localizes to the Golgi in endothelial cells. (A) Immunoblot of sFLT1-HA in concentrated conditioned media or cell lysates of HUVEC transfected with indicated DNA and probed with indicated antibodies. Tubulin loading control. (B) HUVEC immunofluorescence of sFLT1-HA or control and stained for HA, γ-Adaptin (Trans-Golgi), and DAPI (nucleus). Scale bar, 20 μm. Yellow line, line scan. (B’) Line scan of sFLT1-HA and γ-Adaptin fluorescence intensity. (C) HUVEC immunofluorescence with indicated antibodies: FLT1, γ-Adaptin (Trans-Golgi), and DAPI (nucleus). Scale bar, 20 μm. Yellow line, line scan. (C’) Line scan of FLT1 and γ-Adaptin fluorescence intensity. (D) HUVEC density ultracentrifugation subcellular fractionation (see Methods for details). Dashed box, fractions positive for both sFLT1 and STX6 (Trans-Golgi marker). Other markers: Calnexin (endoplasmic reticulum); ZO1 (plasma membrane); EEA1 (vesicles) GAPDH (cytosol). (A-D) n=3 replicates.

### sFLT1 secretion from endothelial cells requires the Golgi, clathrin, and SNAREs

To establish broad requirements for sFLT1 trafficking and secretion in endothelial cells, primary endothelial cells were exposed to pharmacological inhibitors that block trafficking at specific steps of the trafficking process. Brefeldin-A blocks protein transport from the ER to the Golgi, and treatment led to significant reduction of sFLT1 in HUVEC conditioned media, while lysate levels trended towards a slight elevation **(Fig. 3A**, **Supp Fig. 3A,F)**, consistent with constitutive trafficking of sFLT1 through the Golgi and confirming a previous report [2].

**Fig. 3.**
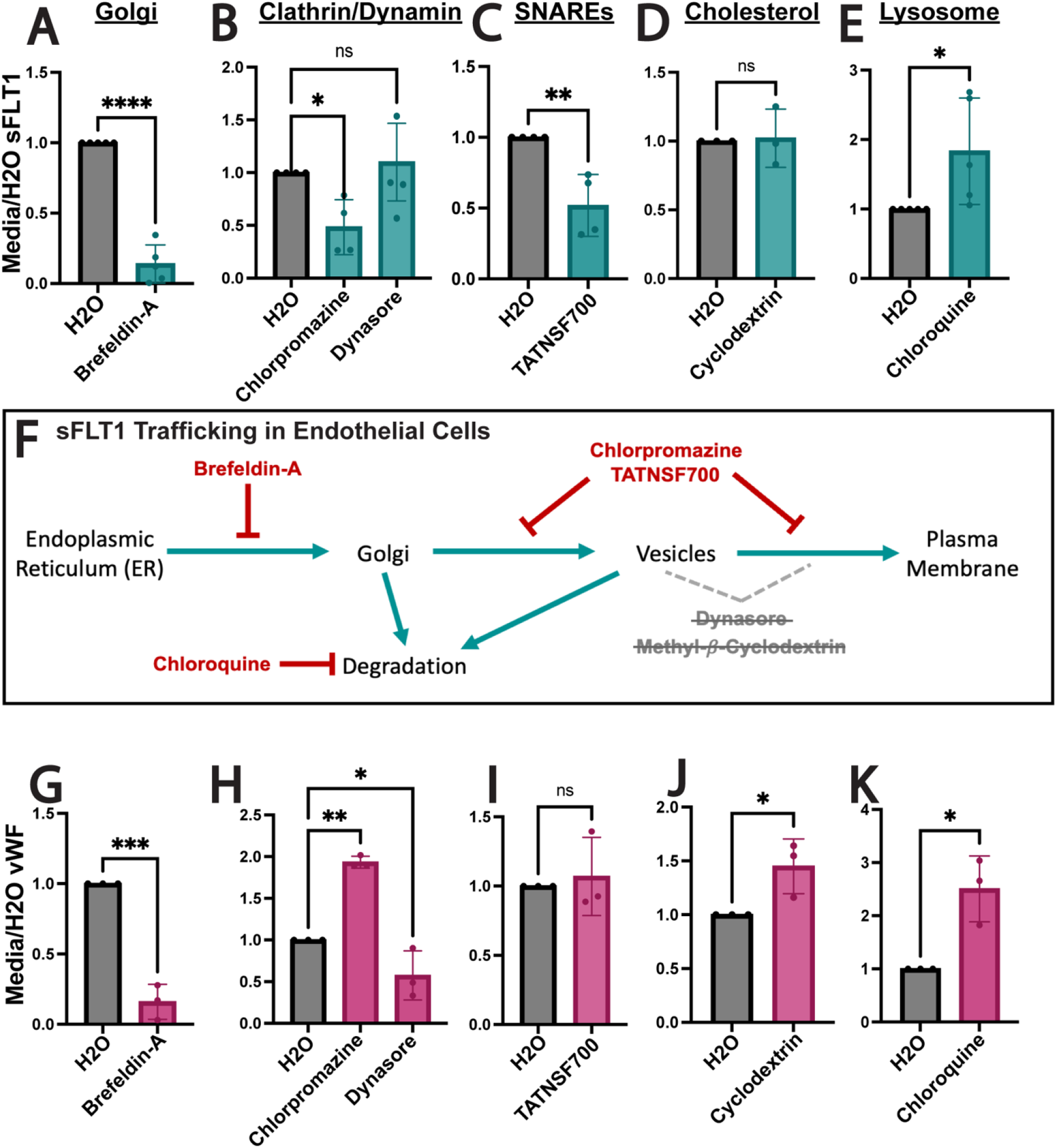
Endothelial sFLT1 trafficking requires the Golgi, clathrin, and SNAREs. (A-E, G-K) Quantification of immunoblots of HUVEC treated as indicated (X-axis) prior to collection of concentrated media or lysates and incubated with FLT1 (A-E) or vWF (G-K) antibody and normalized to vehicle control. Mean +/− SD of experimental replicates shown. Statistics: student’s two-tailed t-test for 2 conditions or one-way ANOVA with pairwise comparison and post-hoc Tukey’s range test for 3 conditions, *P<0.05, **P<0.01, ***P<0.001, ****P<0.0001, ns, not significant. (F) Diagram of sFLT1 protein trafficking steps tested. Red: inhibitors that impacted sFLT1 secretion. Grey with cross-out: inhibitors that did not impact sFLT1 secretion.

Although more commonly known for their involvement in endocytic recycling pathways [68], clathrin and dynamin can also facilitate Golgi-to-vesicle trafficking [69–72]. Dynamin is implicated in constitutive secretion [73]; however, clathrin mainly regulates specialized pathways such as trafficking of Weibel-Palade bodies that leads to secretion of vWF [74]. To determine if sFLT1 constitutive trafficking and secretion are clathrin- or dynamin-dependent, HUVEC were treated with chlorpromazine to block clathrin-mediated or dynasore to block dynamin-mediated trafficking. Chlorpromazine treatment significantly reduced sFLT1 levels in the conditioned media, while lysate levels trended to increase **(Fig. 3B**, **Supp Fig. 3B,G)**, indicating a role for clathrin-coated vesicles in sFLT1 secretion. sFLT1 levels in the media and lysates were unaltered by dynasore; therefore, dynamins are likely not required for sFLT1 secretion **(Fig. 3B**, **Supp Fig. 3B,H)**.

Next, we inhibited trafficking components that promote the release of secreted proteins from the plasma membrane. SNARE complexes are essential for fusion of vesicles with target membranes; therefore, we inhibited SNARE disassembly with the NSF analog peptide, TATNSF700 [75] and found that sFLT1 secretion was reduced while cellular sFLT1 levels were minimally affected **(Fig. 3C**, **Supp Figs. 3C,I)**. Cholesterol-enriched lipid rafts are sites of localized membrane trafficking [76,77], and lipid rafts have been implicated in sFLT1 binding to the surface of podocytes [78]. We assessed the role of lipid rafts in sFLT1 secretion via cholesterol depletion using methyl-*β*-cyclodextrin; however, neither secreted or cell-associated endothelial sFLT1 levels were altered by cholesterol depletion **(Fig. 3D**, **Supp Figs. 3D,J)**.

Finally, endothelial sFLT1 degradation was interrogated with chloroquine treatment to block lysosome-mediated degradation. Both secreted and cell-associated sFLT1 accumulated following chloroquine treatment of endothelial cells, consistent with a lysosomal degradation mechanism for sFLT1 **(Fig. 3E**, **Supp Fig. 3E,K)**. In summary, pharmacological trafficking perturbations showed that endothelial sFLT1 constitutive secretion is dependent on ER to Golgi transport, that vesicle trafficking of sFLT1 from the Golgi to the plasma membrane is SNARE-dependent and unexpectedly dependent on clathrin, and that intracellular sFLT1 is degraded by the lysosome **(Fig. 3F)**.

The requirements for endothelial sFLT1 secretion were then compared to requirements for constitutive secretion of vWF, an endothelial cell protein that is secreted both constitutively and upon stimulation via Weibel-Palade bodies [79,80,48,49]. Constitutive vWF secretion was reduced by brefeldin-A treatment, which blocks ER to Golgi transport, and blockade of lysosomal degradation via chloroquine increased vWF secretion, consistent with previous findings [49,48] and similar to sFLT1 sensitivities **(Fig. 3G,K**; **Supp Fig. 3F,K)**. However, several other manipulations produced different results between constitutive sFLT1 and vWF secretion profiles in endothelial cells. Secreted vWF significantly increased with clathrin blockade via chlorpromazine compared to reduced secretion of sFLT1 **(Fig. 3H**, **Supp Fig. 3G)**. This increase is likely due to a requirement for clathrin in Weibel-Palade body maturation, such that vWF is shunted to the constitutive pathway upon clathrin blockade [74]. In contrast to the sFLT1 secretion profile, vWF secretion was inhibited by dynamin blockade via dynasore treatment, suggesting that dynamins are involved in constitutive vWF trafficking **(Fig. 3H**, **Supp Fig. 3H)**. vWF secretion was unaffected by the blockade of SNAREs with TATNSF700 which reduced sFLT1 secretion **(Fig. 3I**, **Supp Fig. 3I)**, and cholesterol depletion via methyl-*β*-cyclodextrin elevated constitutively secreted vWF levels while not affecting sFLT1 secretion **(Fig. 3J**, **Supp Fig. 3J)**. Thus, although sFLT1 and constitutive vWF secretion both depend on Golgi transport and lysosomal degradation, the divergent sensitivity profiles indicate that sFLT1 and vWF utilize different trafficking pathways to move from the Golgi to the plasma membrane for constitutive secretion from endothelial cells.

### Intracellular sFLT1 is mis-localized following trafficking perturbations

To determine how sFLT1 trafficking perturbations affect intracellular localization, we used immunofluorescence to determine sFLT1-HA and FLT1 reactivity upon pharmacological blockade of endothelial cell trafficking/secretion. Brefeldin-A treatment that blocked the Golgi resulted in mis-localization of both sFLT1-HA and endogenous FLT1 signal from the Golgi to the ER **(Fig. 4A, a-b; 4B, a-b)**, consistent with previous findings [2]. Blockade of clathrin assembly through chlorpromazine treatment increased both sFLT1-HA and FLT1 intensity within the Golgi and in large perinuclear puncta compared to controls **(Fig. 4A, a,c; 4B, a,c)**, consistent with clathrin being required for sFLT1 trafficking from the Golgi to vesicles.

**Fig. 4.**
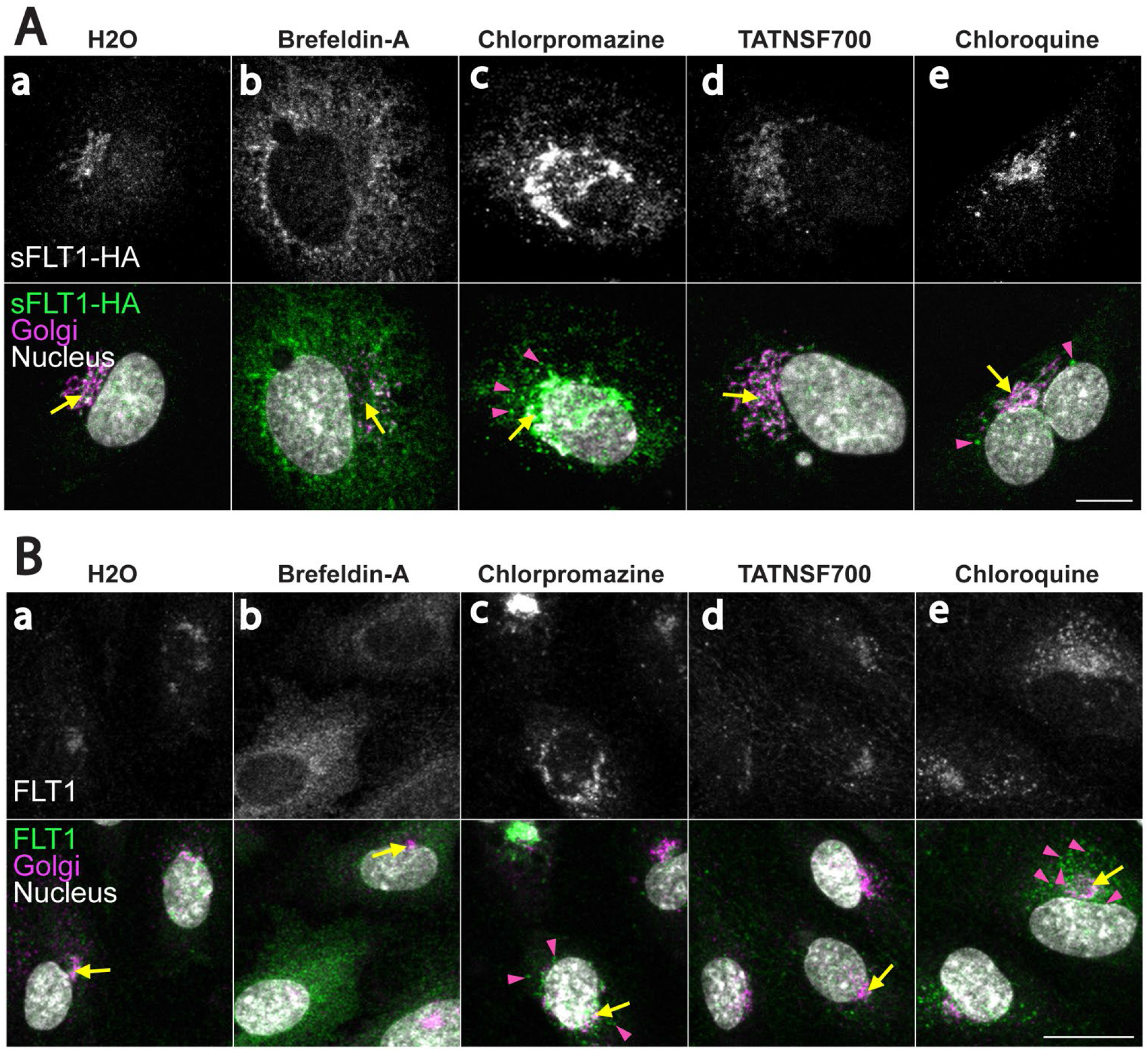
Intracellular sFLT1 is mis-localized following trafficking perturbations. (A, a-e; B, a-e) HUVEC immunofluorescence stained for sFLT1-HA (A, a-e) or FLT1 (B, a-e), GM130 (Golgi), and DAPI (nucleus) after 4 hr of indicated pharmacological inhibitor treatments. n=3 replicates. Scale bar, 10 μm (A), 20 μm (B). Yellow arrows: Golgi-localized sFLT1-HA (A) or FLT1 (B); pink arrowheads: sFLT1-HA (A) or FLT1 (B) localized to puncta.

TATNSF700 did not affect sFLT1-HA or FLT1 localization **(Fig. 4A, a,d; 4B, a,d)**, perhaps because vesicle movement in the cytoplasm is below the sensitivity of detection. Finally, after inhibition of lysosomal degradation with chloroquine treatment, sFLT1-HA and endogenous FLT1 reactivity became more pronounced in large puncta surrounding the Golgi **(Fig. 4A, a,e; 4B, a,e)**, consistent with intracellular accumulation in the absence of degradation. Overall, intracellular sFLT1 localization and sFLT1 secretion had similar sensitivity profiles to the same manipulations that revealed the unexpected requirement for clathrin at the Golgi-to-vesicle step of trafficking of sFLT1 in endothelial cells.

### sFLT1 in polarized vessels and *in vivo* is sensitive to clathrin inhibition

Since trafficking dynamics *in vivo* occur in a 3D environment, we assessed sFLT1 intracellular localization following pharmacological trafficking manipulations in polarized blood vessels. Using a 3D sprouting angiogenesis assay that results in lumenized endothelial cell sprouts polarized in several axes [81,82], FLT1 signal was detected in the Golgi and vesicles of control HUVEC sprouts **(Fig. 5A, a)**, and FLT1 in endothelial cell sprouts displayed similar localization changes following inhibitor treatments as seen in 2D culture. FLT1 became diffuse and ER-localized following brefeldin-A treatment **(Fig. 5A, b)**, while treatment with either chlorpromazine to block clathrin assembly or chloroquine to block lysosomal degradation resulted in FLT1 accumulation in perinuclear puncta **(Fig. 5A, c,d)**. These data are consistent with a requirement for Golgi trafficking, clathrin, and lysosomal degradation in the proper localization of intracellular FLT1 in polarized vessels.

**Fig. 5.**
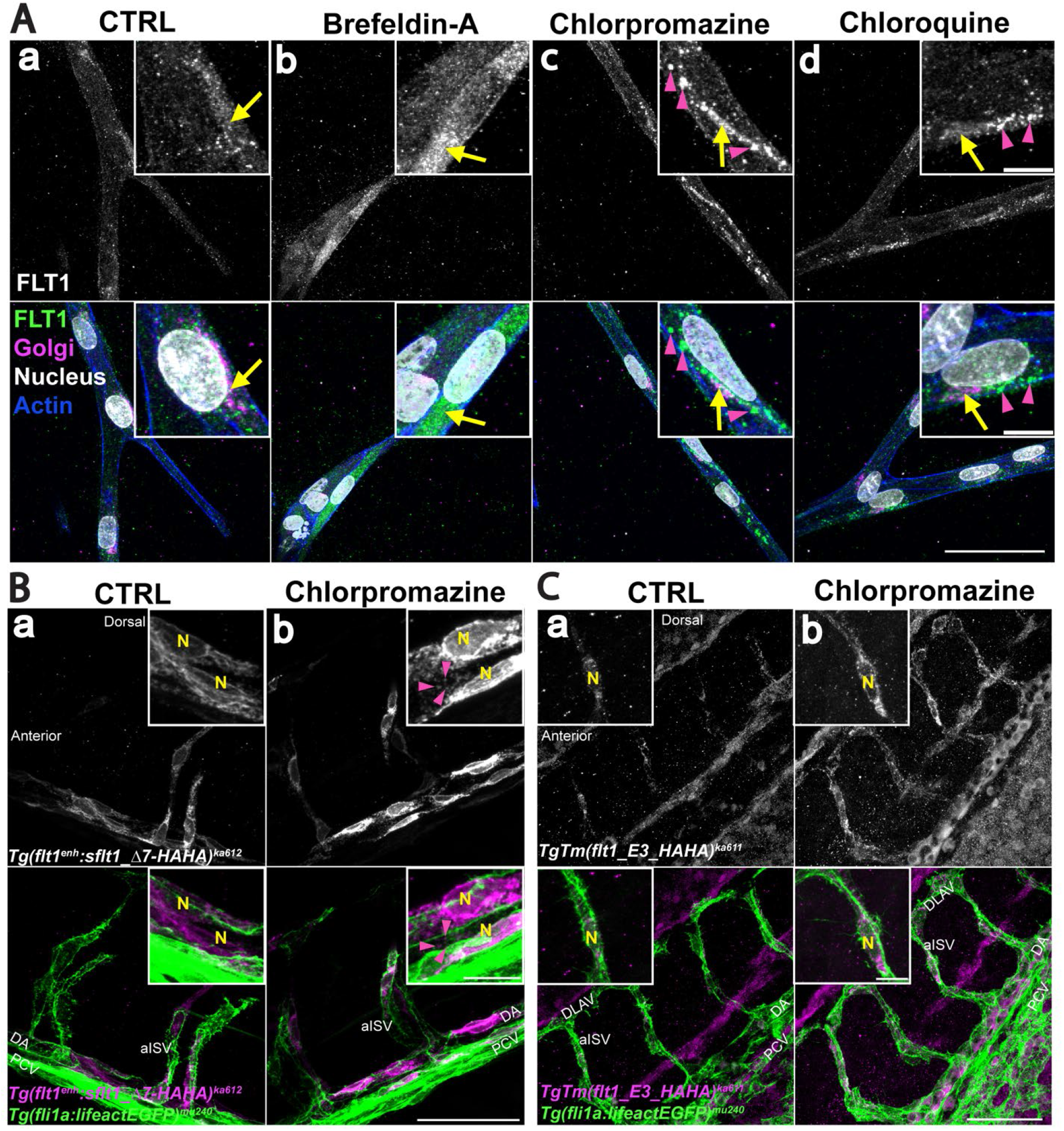
sFLT1 localization in polarized vessels and *in vivo* is sensitive to trafficking perturbations. (A, a-d) Immunofluorescence of day 7 3D HUVEC sprouting assay with indicated inhibitor treatments (18hr) and stained for FLT1, γ-Adaptin (Golgi), DAPI (nucleus), and phalloidin (actin). Scale bar, 50 μm; inset scale bar: 10 μm. n= 3 replicates, 5 beads per replicate. Yellow arrows: Golgi-localized FLT1; pink arrowheads: FLT1 localized to puncta. (B, a-b) Immunofluorescence of 32 hpf *Tg(flt1^enh^:sFLT1_△7-HAHA)^ka612^; Tg(fl1ai:lifeactEGFP)^mu240^* zebrafish embryos labeled with HA antibody following 6 hr of chlorpromazine treatment. N, nucleus; pink arrowheads: HA localized to puncta. Scale bar, 50 μm; inset scale bar: 10 μm. n= 3 replicates. (C, a-b) Representative images of 32 hpf *TgTm(flt1_E3_HAHA)^ka611^; Tg(fl1ai:lifeactEGFP)^mu240^* zebrafish embryos labeled with HA antibody following 6 hr of chlorpromazine treatment. Scale bar, 50 μm; inset scale bar: 10 μm. n= 3 replicates. N, nucleus. DA, Dorsal Aorta. PCV, posterior cardinal vein. aISV, arterial intersegmental vessel. DLAV, dorsal longitudinal anastomotic vessel.

To assess the unique requirement of clathrin for sFLT1 localization in polarized vessels *in vivo*, *Tg(fli1a:LifeAct-GFP)^mu240^* zebrafish embryos were micro-injected with a tagged *sflt1* construct, *flt1^enh^:sflt1_Δ7-HAHA*, at the one-cell stage and analyzed at 32 hours post-fertilization (hpf). Overexpression of wild-type *sflt1* in zebrafish is lethal, so the construct contains a short deletion in the VEGF-binding domain that is predicted to prevent VEGF binding and is tolerated by fish embryos [83]. Mosaic expression of *sflt1_Δ7-HAHA* was detected in the dorsal aorta and arterial intersegmental vessels of the trunk, using an HA antibody **(Fig. 5B, a)**. Under control conditions, HA signal localized to the perinuclear space of endothelial cells. Chlorpromazine treatment to block clathrin resulted in accumulation of HA signal around the nucleus and in puncta of the zebrafish endothelial cells, similar to the pattern seen *in vitro* with these manipulations **(Fig. 5B, b)**.

We next investigated localization of endogenous zebrafish *flt1* using a transgenic line in which a HA-tag was inserted into the *flt1* locus, *TgTm(flt1_E3_HAHA)^ka611^* line [83]. HA antibody staining in the arterial intersegmental vessels of 32 hpf *TgTm(flt1_E3_HAHA); Tg(fli1a:LifeAct-GFP)* embryos was largely perinuclear in controls, and consistent with the HUVEC results and *sflt1_Δ7-HAHA* embryos, HA signal accumulated throughout the cell following chlorpromazine treatment **(Fig. 5C, a,b)**. Overall, sFLT1 localization and sensitivity profiles in 2D cultures were similar in 3D polarized vessels *in vitro* and *in vivo*, and clathrin depletion mis-localized sFLT1 and blocked secretion in 2D culture as well as in polarized vessels, indicating similar requirements for sFLT1 trafficking in physiologically relevant topologies.

### Endothelial sFLT1 secretion requires RAB27a and SNAREs

To define a molecular pathway for sFLT1 trafficking/secretion in endothelial cells, we used siRNA-mediated depletion in primary endothelial cells to target specific trafficking components. Post-Golgi proteins have several potential routes for secretion; some are directly targeted to the plasma membrane, while others fuse with storage granules, recycling endosomes, or other intermediates prior to plasma membrane docking and secretion [84,85,42]. Due to the technical challenges of studying small, fast-moving vesicles, the molecular components of constitutive secretion are not well-defined. However, several RABs and SNAREs that localize to recycling endosomes or storage granules in endothelial cells have been identified [86–88], and the RAB GTPase RAB27a is required to transport and dock Weibel-Palade bodies in endothelial cells for stimulated secretion of vWF [89,90]. We found that RAB27a silencing significantly blocked constitutive sFLT1 secretion without significant reduction of internal sFLT1 levels **(Fig. 6A**, **Supp Fig. 4A,C).** In contrast, depletion of the recycling endosome-localized RABs RAB4a, RAB11a, or RAB8a did not significantly affect sFLT1 secretion or internal sFLT1 levels compared to non-targeting controls **(Fig. 6A**, **Supp. Fig. 4A, D-F)**. Therefore, endothelial cell post-Golgi transport of sFLT1 to the plasma membrane requires RAB27a but not RABs associated with recycling endosomes, suggesting that sFLT1 trafficking requirements overlap with requirements for granule formation and Weibel-Palade body transport.

**Fig. 6.**
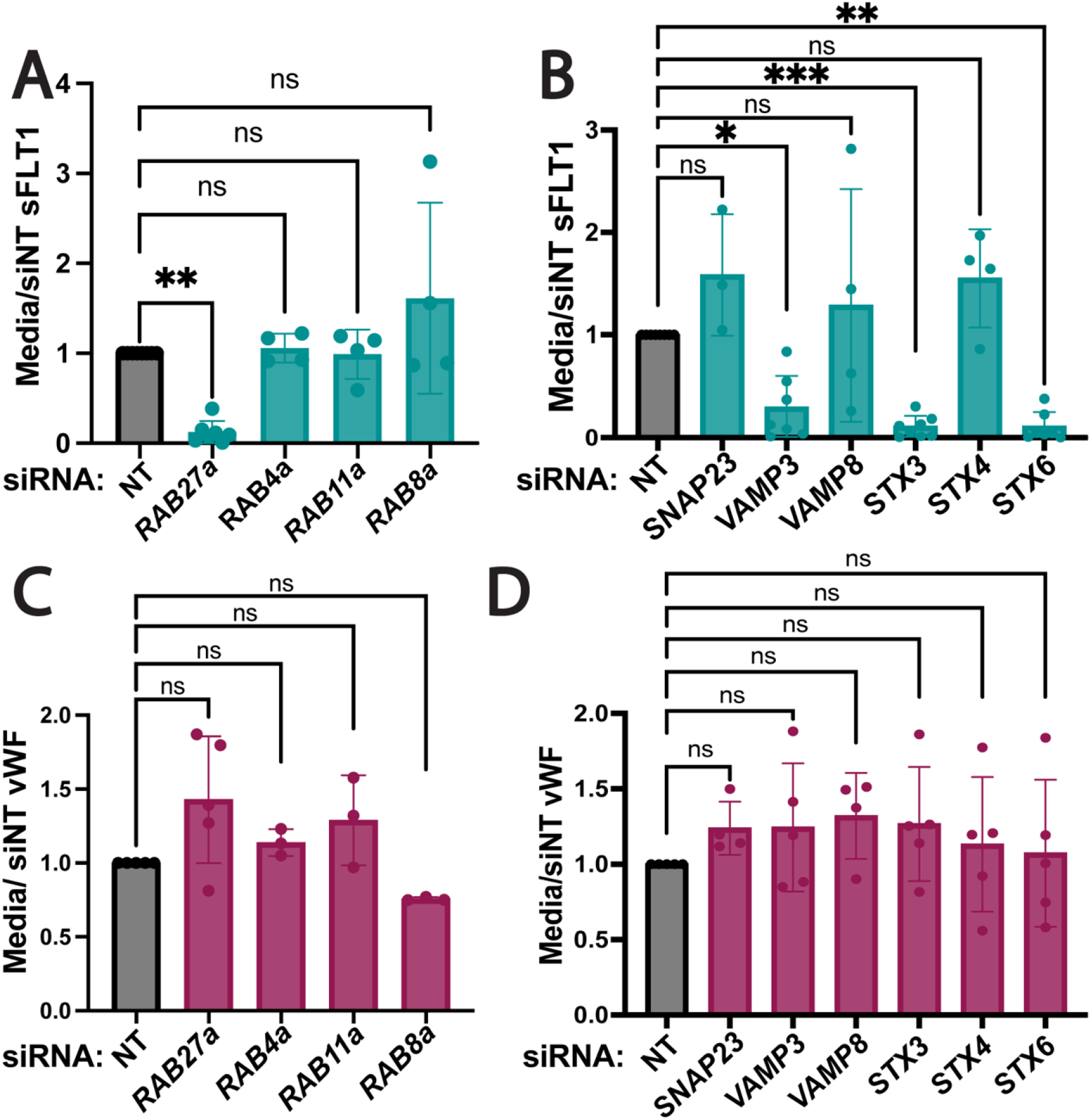
sFLT1 trafficking requires RAB27a and SNAREs. Quantification of sFLT1 (A,B) or vWF (C,D) immunoblot band intensity in 18 hr HUVEC concentrated media following indicated RAB (A,C) or SNARE (B,D) siRNA treatments normalized to siNT. Statistics: Mean +/− SD per experiment. One-way ANOVA with pairwise comparison and post-hoc Tukey’s range test. *P<0.05, **P<0.01, ***P<0.001, ns, not significant.

Fusion of Golgi-derived vesicles with recycling endosomes, storage granules, and the plasma membrane requires SNARE complex formation [39]. The functional SNARE inhibitor, TATNSF700, blocked sFLT1 secretion from endothelial cells; therefore, a panel of endothelial SNAREs was analyzed to determine specificity in sFLT1 trafficking. The SNAREs SNAP23, STX3, STX4, VAMP3, and VAMP8 promote fusion of Weibel-Palade bodies with the plasma membrane for stimulated secretion of vWF in endothelial cells, and STX6 mediates trans-Golgi vesicle trafficking events including localization to clathrin-coated membranes and recycling endosomes [46,91,45,92-94]. Depletion of STX6, STX3 or VAMP3 significantly blocked sFLT1 secretion while si*SNAP23,* si*VAMP8*, and si*STX4* treatment did not affect sFLT1 secretion **(Fig. 6B**, **Supp. Fig. 4G-K)**. Internal sFLT1 levels did not significantly change relative to controls after any SNARE depletions **(Supp. Fig. 4B)**. Together, these data suggest that STX6, STX3, and VAMP3 are required for sFLT1 secretion from endothelial cells. We compared sFLT1 requirements for RAB27a, STX6, STX3, and VAMP3 for endothelial cell secretion to requirements for constitutive vWF secretion, and we found that unlike stimulated release of vWF through Weibel-Palade bodies, constitutive vWF secretion was not significantly changed by depletion of any of the RABs or SNAREs tested compared to controls **(Fig. 6C-D)**, consistent with published work.

### Trans-Golgi trafficking is required for angiogenic sprouting

Trans-Golgi trafficking is the rate-limiting step of constitutive secretion [72]; therefore, we further analyzed the relationship of intracellular sFLT1 and trans-Golgi localized STX6. Immunofluorescence localization in HUVEC revealed that sFLT1-HA and FLT1 colocalized with STX6 at the Golgi under control conditions **(Fig. 7A-B)** and remained Golgi-localized upon STX6 depletion, although FLT1 antibody signal was reduced in EC silenced for STX6 **(Fig. 7C-D).** FLT1 loss or reduction increases angiogenic sprouting *in vitro* and *in vivo* [22,95,57,96,97,35], consistent with its function as a negative regulator of VEGF-A signaling. To test if STX6-mediated trans-Golgi trafficking impacts angiogenic sprouting, we subjected STX6-depleted HUVEC to the 3D sprouting assay. We confirmed that depletion of either total FLT1 (both mFLT1 and sFLT1) or only sFLT1 led to increased endothelial sprouting, with significantly increased sprout numbers and total vessel length **(Fig. 7E-G, I-J)**. STX6 depletion phenocopied the increased sprouting parameters **(Fig. 7H-J)**, consistent with the idea that disrupting the critical Golgi-to-vesicle step of sFLT1 secretion from endothelial cells contributes to angiogenic sprouting defects.

**Fig. 7.**
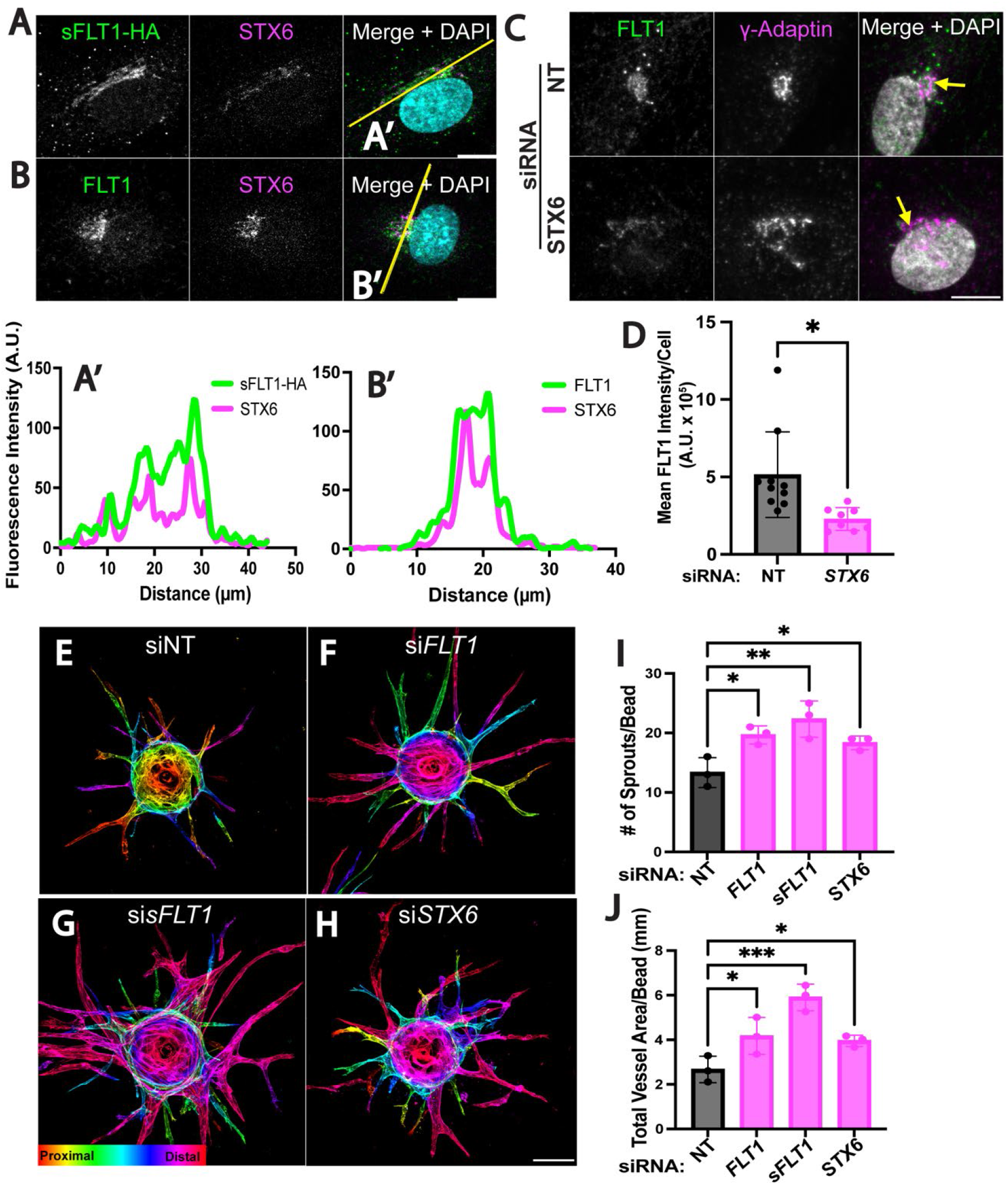
STX6 co-localizes with sFLT1 and regulates angiogenic sprouting. (A,B) HUVEC immunofluorescence imaging of sFLT1-HA (A) or FLT1 (B) and STX6 and DAPI. Scale bar, 20 μm. Yellow line, line scan. (A’-B’) Line scan analysis of sFLT1-HA (A’) or FLT1 (B’) and STX6 fluorescence intensity. (C) Representative immunofluorescence staining of FLT1, γ-adaptin, and DAPI in HUVEC transfected si*STX6* or siNT. Scale bar, 10 μm. (D) Mean FLT1 intensity analysis per cell of HUVEC transfected si*STX6* or siNT. (E-H) Immunofluorescence imaging of 3D HUVEC following indicated siRNA transfections and 4 days of sprouting. Phalloidin staining depth encoded. Scale bar: 100 μm. n= 3 replicates, 3 beads per replicate. (I) Quantification of sprout number/bead. (J) Quantification of the total vessel area/bead (μm). Statistics: (D) Mean +/− SD of FLT1 intensity per nuclei, representative of n= 3 replicates, students two-tailed t-test. *P<0.05; (I,J) Mean +/− SD of 3 beads per experiment, n= 3 replicates, one-way ANOVA with pairwise comparison and post-hoc Tukey’s range test. *P<0.05, **P<0.01, ***P<0.001.

### Golgi-to-vesicle sorting of sFLT1 is mediated by AP1

We further evaluated the role of Golgi trafficking in sFLT1 secretion by depleting Golgi-localized trafficking proteins or their plasma membrane-localized counterparts and assessing effects on sFLT1 secretion and intracellular localization. ARF1, a downstream target of brefeldin-A, is required at the Golgi to recruit COPI coated vesicle proteins and clathrin adaptors such as AP1 [98]. In contrast, ARF6 assists in rearranging actin near the plasma membrane [98]. HUVEC were treated with siRNAs targeting ARF1 or ARF6, and si*ARF1* blocked accumulation of sFLT1 in the media compared to controls without affecting sFLT1 levels internally **(Fig. 8A**, **Supp. Fig. 5A, D-E),** whereas ARF6 depletion did not alter secreted or intracellular sFLT1 levels **(Fig. 8B**, **Supp. Fig. 4B,E)**. These findings indicate that Golgi-localized ARF1 regulates sFLT1 secretion while peripheral ARF6 does not affect secretion.

**Fig. 8.**
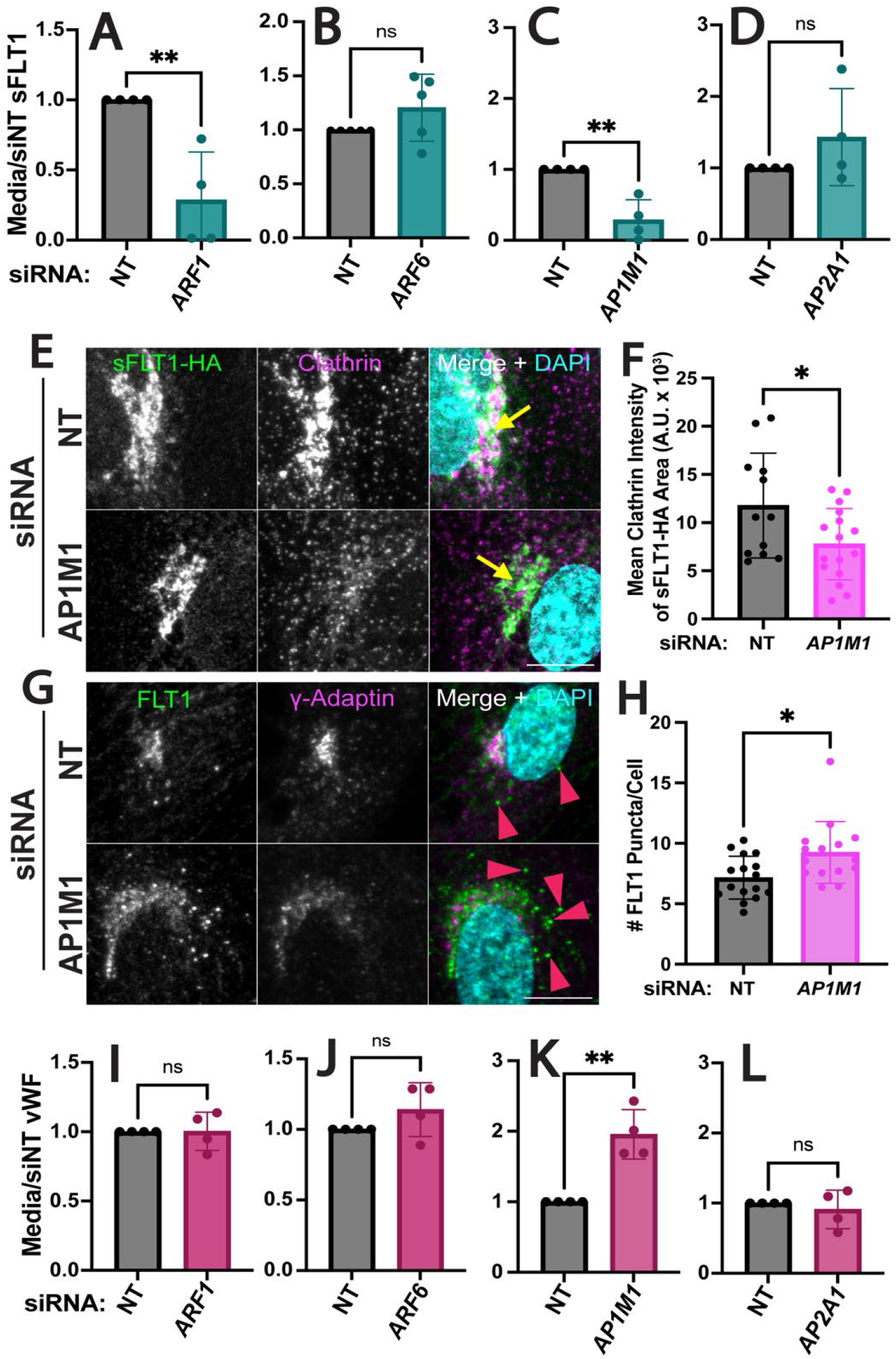
Golgi trafficking of sFLT1 is clathrin-mediated. (A-D) Quantification of sFLT1 immunoblot band intensity in HUVEC conditioned media with indicated siRNA treatments normalized to siNT. (E,G) Immunofluorescence staining of sFLT1-HA and clathrin (E) or FLT1 and γ-adaptin (G) and DAPI in HUVEC transfected with si*AP1M1* or siNT. Yellow arrows: Golgi-localized sFLT1-HA; pink arrowheads: FLT1 accumulation in puncta. Representative of n=3 replicates. Scale bar, 10 μm. (F) Quantification of clathrin intensity localized to sFLT1-HA area, n= 3 replicates, (H) Quantification of FLT1 puncta per cell, n=3 replicates. (I-L) Quantification of vWF immunoblot band intensity in HUVEC conditioned media with indicated siRNA treatments normalized to siNT. Statistics: Mean +/− SD of experimental replicates shown. students two-tailed t-test. *P<0.05, **P<0.01, ns, not significant.

Finally, to determine whether the unexpected clathrin requirement for sFLT1 secretion was linked to the Golgi, we depleted the *μ*1 subunit of the Golgi-localized adaptor protein, AP1 (si*AP1M1*), or the *α*1 subunit of plasma membrane-localized AP2 (si*AP2A1*) [71], as complexes with each subunit are predicted to promote clathrin-dependent trafficking at different cellular locations. sFLT1 secretion was inhibited by si*AP1M1* but not si*AP2A1*, indicating that clathrin-dependent sFLT1-trafficking likely occurs at the Golgi but not the plasma membrane **(Fig. 8C-D**, **Supp. Fig. 5F-G)**. As described earlier, sFLT1-HA and FLT1 colocalize with the gamma subunit of AP1, γ-Adaptin, in the Golgi; internal sFLT1 levels were reduced after si*AP1M1* but not si*AP2A1* treatment compared to controls **(Supp. Fig. 5C, F-G)**. Immunofluorescence imaging of sFLT1-HA and FLT1 revealed that AP1M1 depletion reduced colocalization of clathrin with Golgi-localized sFLT1-HA **(Fig. 8E-F)**, while FLT1 signal increased in large perinuclear puncta **(Fig. 8G-H)**. Therefore, sFLT1 is likely sorted into a clathrin-dependent compartment at or near the Golgi that is required to target sFLT1 for secretion.

Together, these findings establish a requirement for ARF1 and AP1 at the Golgi for proper trafficking and secretion of constitutive sFLT1. To determine if constitutive vWF also exhibits these requirements in endothelial cells, we interrogated constitutive vWF secretion in parallel. Neither ARF1 or ARF6 depletion affected accumulation of constitutive vWF in the media **(Fig. 8I-J)**. Constitutive vWF secretion significantly increased following AP1M1 depletion and did not change with AP2A1 depletion **(Fig. 8K-L)**, consistent with published work [74]. Therefore, sFLT1 trafficking in endothelial cells diverges from constitutive vWF trafficking in the same cells at or near the Golgi.

## DISCUSSION

Here we define a unique trafficking pathway for endothelial cell secretion of sFLT1 **(Fig. 9)**, a critical regulator of VEGF-A signaling involved in blood vessel formation and vascular pathologies. Constitutive sFLT1 secretion utilizes STX6, ARF1, and AP1-dependent clathrin at or near the Golgi, likely for sorting from an intermediate compartment for trafficking to the plasma membrane via RAB27a and specific SNAREs. Blockade or depletion of trafficking/secretion components led to sFLT1 mis-localization within endothelial cells, in polarized 3D vessels, and *in vivo* in zebrafish. Thus, our findings highlight the importance of sFLT1 trafficking and secretion in the regulation of blood vessel formation and provide new targets for regulation of the pathway.

**Fig. 9.**
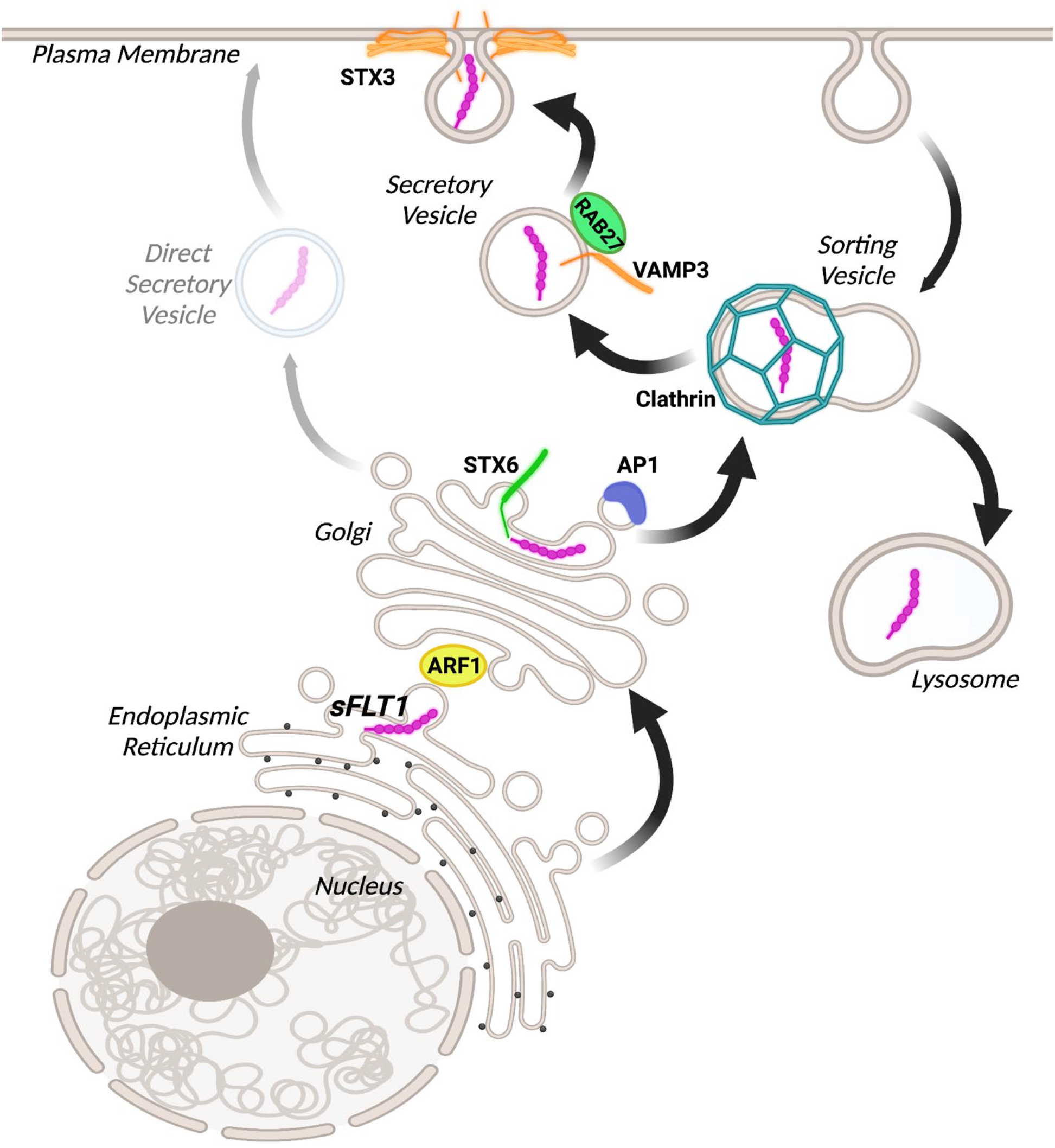
Model for sFLT1 trafficking and secretion from endothelial cells. The data is consistent with a model whereby sFLT1 is trafficked between the endoplasmic reticulum and Golgi by ARF1, the Golgi and a clathrin-coated sorting compartment by STX6 and AP1, and finally through constitutive vesicles to the plasma membrane by RAB27a, VAMP3, and STX3 or to the lysosome for degradation. An additional direct secretory pathway from the Golgi may be utilized although it does not compensate for the loss of the depicted trafficking components. Figure made with Biorender.com.

We used computational modeling to predict a sFLT1 secretion rate of approximately 30,000 molecules/cell/hr, which is well in excess of what is likely needed to create significant reservoirs of interstitial sFLT1 in tissues [99]. Notably, our estimated endothelial sFLT1 secretion rate is also higher than the secretion rate of the ligand VEGF-A from stromal cells (~250-1250 molecules/cell/hr) [59]. This implies that in most well-vascularized tissues, interstitial sFLT1 levels are predicted to exceed those of VEGF-A, consistent with a role for sFLT1 in modulating VEGF-A signaling and regulating angiogenesis. Our mechanistic model parameters are consistent with sFLT1 secretion time courses across 3 independent studies, suggesting that this model will be useful in the future to determine how individual processes contribute to sFLT1 secretion and to simulate mechanistic interventions.

Three main routes target proteins for secretion from the Golgi: small unregulated vesicles, large secretory granules that are released upon stimulus, or sorting into an intermediate compartment prior to secretion [85,84,42]. Endothelial cell sFLT1 secretion does not require a stimulus [51–53], and our data shows that sFLT1 is constitutively secreted utilizing clathrin, suggesting that sFLT1 is sorted into an intermediate compartment prior to trafficking to the cell surface. Although we cannot rule out that some sFLT1 is also secreted via small unregulated vesicles, disruption of the clathrin-mediated pathway via pharmacological blockade or depletion of clathrin-associated components did not lead to rescue of sFLT1 secretion by alternate trafficking routes, indicating that the major endothelial cell sFLT1 trafficking pathway utilizes a clathrin-dependent intermediate.

Although clathrin-mediated trafficking is linked to recycling via endosomes from the plasma membrane [68], we found no evidence that sFLT1 secretion utilizes traditional surface-endosomal recycling trafficking proteins (i.e. ARF6, RAB4, RAB11, and RAB8 or the plasma-membrane clathrin adaptor AP2), consistent with the idea that secreted proteins are usually not recycled from the cell surface. These findings instead suggest that sFLT1 primarily utilizes a poorly characterized intermediate sorting pathway to exit endothelial cells. AP1-mediated clathrin assembly is required for the maturation of immature Golgi-derived secretory granules in many cell types, including endothelial cells [100,74], and AP1 works with clathrin to sort uncondensed proteins for trafficking as they condense into secretory granules, likely to target lysosomes or the plasma membrane [101]. Proteins trafficked via this mechanism include lysosomal enzymes that interact with mannose-6-phosphate receptors for lysosomal sorting and proinsulin that is sorted from condensed insulin into immature granules for constitutive secretion [102,101,103].

Evidence that sFLT1 trafficking utilizes a clathrin-dependent intermediate compartment is also supported by our finding of a requirement for STX6 for sFLT1 secretion, since STX6 colocalizes with AP1/clathrin on immature granules and regulates Golgi-to-vesicle transport [104,105,102]. Moreover, our finding that sFLT1 and STX6 colocalize through cell fractionation and immunofluorescence imaging suggests a similar requirement for STX6 in the trafficking of sFLT1 from the Golgi. Endothelial sFLT1 localizes to large puncta and is degraded by the lysosome; thus, our findings are consistent with a model whereby newly synthesized sFLT1 utilizes a post-Golgi immature granule-like compartment to move to the plasma membrane for secretion or to the lysosome for degradation **(Fig. 9)**. Blockade of clathrin assembly in sprouting endothelial cells *in vitro* and in zebrafish revealed sFLT1 mis-localization as seen in 2D culture, and depletion of STX6 phenocopied FLT1 depletion in sprouting parameters, leading to excess sprouting and branching, consistent with the idea that Golgi trafficking and secretion of sFLT1 is operative in physiologically relevant settings.

We compared endothelial cell sFLT1 trafficking/secretion requirements to those of vWF, as the stimulated endothelial vWF trafficking pathway is well described [48,49,106,92]. Interestingly, despite being constitutively secreted like unstimulated vWF, sFLT1 utilizes some trafficking components also used for stimulated vWF secretion via Weibel-Palade bodies. Our data revealing a requirement for AP1-mediated clathrin assembly in constitutive sFLT1 secretion is similar to requirements for Weibel-Palade body trafficking and maturation [74], and these requirements are not shared by constitutive vWF secretion assayed in parallel here. Additionally, we show that RAB27a and the specific SNAREs VAMP3 and STX3 are required for constitutive sFLT1 secretion, similar to the requirements for histamine-responsive Weibel-Palade body release but not constitutive vWF secretion [44]. However, sFLT1 is not detectable in mature Weibel-Palade bodies [50], and unlike vWF secreted by Weibel-Palade bodies, sFLT1 secretion has a requirement for the SNARE STX6 but not SNAP23, STX4, or VAMP8. Thus, some trafficking components appear to be shared while others differ between constitutive and stimulated pathways for secretion from endothelial cells, setting up a unique “hybrid” clathrin-dependent trafficking pathway for endothelial sFLT1 secretion.

sFLT1 secretion from endothelial cells is critical to its regulation of VEGF signaling, and our findings reveal novel aspects of sFLT1 trafficking and secretion that define a unique pathway for sFLT1 constitutive trafficking/secretion from endothelial cells. The importance of secreted sFLT1 in vascular function is highlighted by recent work showing that vascular decline is linked to endothelial cell aging and reduced microvascular density, and that these changes are in turn promoted by reduced VEGF signaling resulting from elevated sFLT1 serum levels [28]. It is possible that trafficking/secretion of the soluble FLT1 isoform contributes to increased serum levels of sFLT1 and age-related endothelial cell dysfunction, along with changes in alternative splicing of FLT1 described in the study. Since vascular decline is thought to be a major driver of numerous age-related changes in metabolism and inflammatory responses, the regulation of sFLT1 secretion likely has a central role in maintaining homeostasis throughout life.

## Materials and Methods

### Cell Culture

Human umbilical vein endothelial cells (HUVEC) (Lonza, #C2519A, Lot # 0000661173 and 0000704189) were cultured in EBM-2 media supplemented with a bullet kit (EGM-2) (Lonza, #CC-1362) and 1x antibiotic-antimycotic (Gibco). HUVEC were used from passages 3-6. Normal human lung fibroblasts (NHLF) (Lonza, #CC-2512) were cultured in DMEM (Gibco, #11965118) supplemented with 10% fetal bovine serum (FBS) and 1x antibiotic-antimycotic (Gibco) and used at passages 4-10. HEK293T/17 (ATCC #CRL-11268) were maintained in DMEM supplemented with 10% FBS and 1x antibiotic-antimycotic and used at passages 6-12. All cells were maintained at 37°C and 5% CO_2_. HUVEC, NHLF, and HEK193T/17 cells were certified mycoplasma-free by the UNC Tissue Culture Facility.

### Inhibitor Treatments

For secretion experiments, HUVEC were grown to 80% confluency, washed with PBS, then incubated in Fibroblast Basal Media (Lonza, #CC-3131):EGM-2 (1:1, FBM/EGM) containing pharmacological inhibitors (**Supplementary Table S6**) or vehicle controls overnight at 37°C before protein collection. For trafficking experiments, HUVEC were allowed to adhere overnight at 37°C, then FBM/EGM with inhibitors or controls was added for 4 hr at 37°C before fixation.

### Plasmid Expression

The human sFLT1-HA plasmid construct created through Gibson cloning of human sFLT1 cDNA into a pcDNA3.1 vector containing a C-terminal HA tag (pcDNA3-ALK2-HA, Addgene 80870). Sequencing validation was performed through Genewiz. HUVEC were grown to 95% confluency, trypsinized in 0.05% Trypsin-EDTA (Gibco, #25300-054) at 37°C for 3 min, then collected in 1:1 sterile PBS/new born calf serum (NBCS) (Gibco, #16010-159). Cells were pelleted at 1000 rpm for 5 min and resuspended in 100 μL nucleofector solution (Lonza, #VPB-1002) before adding 2.5 μg of plasmid. The suspension was electroporated into cells using the Amaxa Nucleofector System (D-005 program), and cells were then resuspended and seeded in fresh EGM-2. The cells were left to adhere for 5 hr at 37°C and 5% CO_2_ before adding secretion media (1:1 FBM/EGM) to allow for overnight recovery and to remove dead cells. 24 hr after electroporation, the samples were either fixed with 4% PFA for IF or collected for western analysis.

For expression in HEK293T/17, cells were grown to 75-80% confluency and incubated with plasmid diluted in OptiMem (Gibco, #11058021) and Lipofectamine 3000 + P3000 reagent (ThermoFisher, #L3000015) according to manufacturer’s protocol for 24 hr at 37°C. Cells were rinsed in DMEM + 10% FBS, pelleted and resuspended in DMEM +10% FBS or secretion media (1:1 FBM/EGM) and added to fibronectin-coated culture slides for staining or plates for protein collection. 48 hr post-transfection, media was collected and cells were either fixed with 4% PFA for immunofluorescence or collected for Western blot analysis.

### siRNA Transfection

HUVEC were grown to 75-80% confluency and incubated with siRNAs (**Supplementary Table S7**) diluted in OptiMem (Gibco, #11058021) and Lipofectamine 3000 (ThermoFisher, #L3000015) according to manufacturer’s protocol for 24 hr at 37°C. Cells were pelleted, resuspended in EGM-2, and added to fibronectin-coated culture slides for staining or plates for protein collection. 48 hr post-transfection, secretion media (1:1 FBM/EGM) was added. 72 hr post-transfection, media was collected and cells were either fixed with 4% PFA for IF or collected for western analysis.

### Protein Collection and Western Blots

Conditioned media was collected [48], then cells were lysed in 100-200 μL RIPA buffer containing 1X protease-phosphatase inhibitor cocktail (Cell Signaling, #5872S), as previously described [95]. Conditioned media was centrifuged for 5 min at 2500 g at 4°C to pellet remaining cells, then concentrated (Amicon, #UFC803024) per manufacturer’s protocol and centrifuged for 10 min at 3200 g to 250-500 μL volume, prior to addition of protease-phosphatase inhibitor and sample loading buffer containing 10% DTT. Western blot analysis was adapted from a previous protocol [107]. Briefly, 10-50 μg protein was separated by SDS-PAGE and transferred to PVDF membrane. Membranes were blocked for 1 hr with OneBlock Western-CL Blocking Buffer (Genesee), and incubated with primary antibodies diluted in OneBlock (**Supplementary Table S8**) at 4°C overnight. Membranes were washed 3X in PBST (PBS + 0.1% Tween-20) before adding HRP-conjugated secondary antibodies (**Supplementary Table S9**) in OneBlock and incubating for 1 hr at RT. After 3X wash in PBST, Immobilon Forte substrate (Millipore, #WBLUF0100) was added for 1 min. The membranes were then imaged using the ChemiDoc XRS with Chemi High Resolution setting. Restore Western Blot Stripping Buffer (ThermoFisher, 21059) was used for reprobing.

### Cell Fractionation

The cell fractionation protocol was adapted from the OptiPrep Application Sheet (S24). Briefly, HUVEC were grown to confluency, rinsed 2X in PBS, 1X in homogenization media (0.25M sucrose, 1mM EDTA, 10mM Hepes, pH 7.4), and scraped off into homogenization media + 1x protease-phosphatase inhibitor cocktail (Cell Signaling, 5872S). Cells were passed through a 27-gauge needle 15X to remove nuclei, then the homogenate was centrifuged at 1500 g for 10 min at 4°C. The cleared supernatant was added directly to a 2.5-30% iodixanol (OptiPrep, Millipore #D1556) gradient. Solutions of 2.5, 5, 7.5, 10. 12.5, 15, 17.5, 20, or 30% (w/v) iodixanol were prepared with appropriate volumes of homogenization media and 50% iodixanol working solution. The step gradient was formed by bottom loading 13 mL UltraClear tubes (Beckman Coulter, #344059) with a long metal cannula with 800 μL 2.5%, 1.6 mL 5%, 1.6 mL 7.5%, 1.6 mL 10%, 400 μL 12.5%, 1.6 mL 15%, 400 μL 17.5%, 400 μL 20%, and 400 μL 30% iodixanol. The gradients were loaded into a prechilled Beckman SW 41Ti swinging-bucket rotor, then centrifuged at 200,000 g (40,291 rpm) for 2.5 hr; decelerated from 2000 rpm without the brake in a Beckman Coulter ultracentrifuge. 400 μL fractions were carefully collected from the tubes by upward displacement and 5X sample loading buffer containing 10% DTT was added to all samples before boiling for 5 min. Samples were stored at −20°C.

### HUVEC Immunofluorescence Staining

Falcon culture slides (Fisher, #354104) were coated with 5 μg/mL fibronectin for 45 min at RT before seeding 150,000 cells/well and allowing overnight recovery. Following treatments, cells were rinsed with PBS, fixed in 4% PFA for 10 min, then rinsed with PBS and permeabilized with 0.1% TritonX-100 for 10 min at RT. Cells were rinsed with PBS then blocked for 1 hr at RT in blocking solution (5% NBCS +antibiotics + 0.01% Sodium Azide in PBS). Primary antibodies were diluted in blocking solution (**Supplementary Table S8**) and incubation was at 4°C for 48 hr. Cells were washed with PBS 3X before adding Alexa-Fluor-conjugated secondary antibodies **(Supplementary Table S9)**, DAPI, and Alexa-Fluor-conjugated phalloidin in blocking solution and incubating for 3 hr at RT in the dark. Cells were washed with PBS 3X, chambers were removed, and coverslips mounted using Prolong Diamond Antifade mounting media and sealed with nail polish after sitting overnight.

Unless otherwise stated, confocal images were acquired with an Olympus confocal laser scanning microscope and camera (Fluoview FV3000, IX83) and a UPlanSApo 60x oil-immersion objective (NA 1.40) with 1024×1024 resolution and 2x zoom. Images were acquired with the Olympus Fluoview FV31S-SW software and all image analysis, including Z-stack compression, was performed in ImageJ [108,109]. Any adjustments to brightness and contrast were performed evenly for images in an experiment.

### 3D Angiogenesis Sprouting Model

The 3D sprouting angiogenesis assay was performed as previously described [57,81]. Briefly, control HUVEC or HUVEC 48 hr post-siRNA treatment were incubated with cytodex 3 microcarrier beads (17048501, GE Healthcare Life Sciences) overnight then resuspended in 2.2 mg/ml fibrinogen (820224, Fisher) plus aprotinin (A3428, Sigma) and embedded in a fibrin matrix by combining 7 μl of 50U/ml thrombin (T7201-500UN, Sigma) with 500 μl of bead/fibrinogen solution in a 24-well glass-bottomed plate (662892, Grenier Bio) that was incubated for 30 min at RT then 30 min at 37°C. EGM-2 and normal human lung fibroblasts (NHLF, CC2512, Lonza) at a concentration of 2×10^5^ cells/ml was added to each well, and incubation was at 37°C. On day 7 following 18 hr of inhibitor treatment or day 4 following siRNA treatment, fibroblasts were removed via trypsin treatment (5X-trypsin for 7 min at 37°C), and samples were fixed in 4% PFA for 15 min at RT. Samples were permeabilized with 0.5% Triton X-100 in PBS for 1 hr at RT. After rinsing in PBS, samples were blocked in: 5% FBS, 1% donkey serum, 1% BSA (A4503, Sigma), 0.3% Triton X-100 (T8787, Sigma) overnight at 4°C. Samples were rinsed 3X in PBS, then anti-FLT1 (1:500, RnD) and anti-γ-Adaptin (1:500, abcam) antibodies in antibody solution (5% FBS, 1% BSA, 0.1% Triton X-100) were added for 48 hr at 4°C. Samples were rinsed 3X 10 min in 0.5% Tween-20 in PBS then washed overnight at 4°C in 0.5% Tween-20. Samples were rinsed 3X in PBS, then donkey anti-goat AlexaFluor488 (1:1000, ThermoFisher), donkey anti-rabbit AlexaFluor594 (1:1000, ThermoFisher), DAPI (0.3μM, 10236276001, Sigma) and AlexaFluor647 Phalloidin (1:500, Life Technologies) in blocking solution were added to the wells, and incubation was overnight at 4°C prior to rinsing 6X in 0.5% Tween in PBS.

For FLT1 localization, images were acquired in the Z-plane using a UPlanSApo 60x oil-immersion objective (NA 1.40). For whole bead analysis, images were acquired in the Z-plane using a UPlanSApo 20x oil-immersion objective (NA 0.58) and processed in ImageJ. The phalloidin channel was used to create a temporal color code relative to the Z-distance to distinguish overlapping sprouts. The number of sprouts per bead was calculated using the multi-point analysis ImageJ tool. Total vessel area was measured by tracing each vessel from the edge of the bead to the tip of the sprout. The tracing was selected and converted to a mask and skeletonized. The sum of all the branches measured was calculated using the AnalyzeSkeleton plugin, and the pixels were converted to μm.

### Zebrafish

Zebrafish (*Danio rerio*) were housed in an institutional animal care and use committee (IACUC)-approved facility and maintained as previously described [110]. *Tg(fli1a:lifeactEGFP)^mu240^* was received through ZFIN [111]. The *TgTm(flt1_E3_HAHA)^ka611^* line and the *flt1^enh^:sflt1_Δ7-HAHA* Tol2 expression construct were generated as described [83]. As previously described [83,112], 25 ng/uL^−1^ *flt1^enh^:sflt1_Δ7-HAHA* and 25 ng/uL^−1^ Tol2 transposase mRNA were injected into one-cell stage *Tg(fli1a:lifeactEGFP)^mu240^* embryos to generate mosaic expression of tagged sflt1. Tol2 transposase mRNA was generated using the sp6 mMessage mMachine synthesis kit (AM1340, Thermo Fisher). At 26 hpf, *TgTm(flt1_E3_HAHA*)^ka611^ or *Tg(flt1^enh^:sflt1_Δ7-HAHA)* embryos were sorted for lifeactEGFP+ vasculature, dechorinated, and incubated in E3 buffer containing 100 μM Chlorpromazine or H2O vehicle control for 6 hrs at 28.5°C. After treatments, embryos were fixed in ice cold 4% PFA for 2 hr at RT or overnight at 4°C, rinsed 3X with PBS, then permeabilized with 0.1% Tween 20+ 0.5% Triton-X in PBS (PBST-TX) for 1 hr at RT shaking. Embryos were blocked in PBST-TX+ 1% BSA+ 5% goat serum+ 0.01% Sodium Azide for 2 hr at RT shaking. Primary antibodies (**Supplementary Table S8**) were diluted in blocking solution and incubation was overnight at 4°C shaking. Embryos were washed PBST 3X 30 min each at RT followed by overnight wash at 4°C. Secondary antibody (**Supplementary Table S9**) was diluted in blocking solution and incubated for 3 hr at RT in the dark. Embryos were washed 3X with PBST for 30 min at RT, incubated overnight in PBST, then placed onto glass slides for mounting. Heads were removed below the yolk sac with a razor and trunks were mounted in Prolong Diamond Antifade. The slides were cured overnight at RT in the dark before sealing with nail polish.

### Mechanistic computational model of sFLT1 secretion

To quantitatively characterize sFLT1 secretion kinetics, we built a mechanistic model based on two delay differential equations tracking intracellular sFLT1 protein levels (Si) and extracellular sFLT1 protein levels (Sx). The model includes four processes – sFLT1 production, secretion, intracellular degradation, and extracellular degradation (**Fig 1G**; **Supplementary Tables S1-S4**). We parameterized the model using two independent previously published sFLT1 secretion time course data sets from experiments using HUVEC: a pulse-chase experiment tracking fold change in both intracellular and extracellular sFLT1 [2]; and an experiment measuring absolute sFLT1 concentration in conditioned media [1]. The resulting parameters (**Supplementary Table S3**) are also a good fit for the measurements of secreted sFLT1 over time in our study **(Fig 1F)**.

### Data Quantification

Western blot band relative intensity was determined using densitometry analysis [113]. Images were acquired within the linear range using the ChemiDoc XRS with Chemi High Resolution setting. The images were exported as TIFFs and analyzed using the ImageJ Gels tool. An ROI was created around the first lane and copied to each additional lane of the gel. The intensity profiles of each lane was plotted, and a line was drawn to subtract background signal. The pixel intensity of each lane was measured and compared relative to the control lane. Tubulin or GAPDH was used for loading control normalization.

To analyze the spatial organization of sFLT1-HA, FLT1, γ-Adaptin, and STX6 within the Golgi, a parallel line to the longest Golgi axis was drawn in ImageJ. Single plane intensity data of each channel was fitted to a Gaussian distribution to locate the peak of each channel and was then plotted as a function of distance [114].

To determine the overlapping clathrin intensity with sFLT1-HA, single plane images were analyzed in ImageJ. The sFLT1-HA channel was fitted to a Gaussian distribution to obtain the peak intensity region and then a threshold was set to create a mask. The mask was applied to the clathrin channel and average intensity measurements for clathrin within the sFLT1-HA area were acquired. Each sFLT1-HA positive cell per condition across 3 independent experiments was graphed using Prism9 software.

For FLT1 puncta analysis, images with FLT1, a Golgi marker, and DAPI were imported into ImageJ. The Golgi channel was fitted to a Gaussian distribution then thresholded to create a mask. The masked region was then subtracted from the FLT1 channel using the Image Calculator function. The Analyze Particles function was used to count puncta within the size range of 0.2-1 μm. The DAPI channel was used to count the number of nuclei per frame. To obtain the average number of FLT1 puncta per cell, the number of puncta per frame was divided by the number of nuclei. The number of FLT1 puncta per cell from 3 images per 3 replicates was graphed using Prism9 software. For images processed in ImageJ, any changes in brightness and contrast were identical between samples meant for comparison.

### Statistics

Student’s two-tailed t test was used to determine the statistical significance in experiments with two groups. One-way ANOVA with Tukey’s multiple comparisons test was used to determine statistical significance for experiments with 3 or more groups. Error bars represent the mean ± standard deviation of n ≥ 3 independent experiments. Statistical tests and graphs were made using the Prism9 software (GraphPad Software) and R Statistical Software (v 4.0.5, R Core Team, 2021) with the tidyverse package [115].

## Data availability

All data in support of the findings of this work can be found within the article and its Supplementary Information, and from the corresponding author on reasonable request.

## ACKNOWLEGMENTS

We thank Bautch lab members for constructive discussions and Dr. Patrick Brennwald for advice. We thank the UNC Aquaculture Facility, Michelle Altemara, and Michaelanthony Gore for zebrafish husbandry and support, Wendy Salmon at the UNC Hooker Imaging Core for quantitative imaging analysis advice, and Dr. Stephen Rogers for assistance with ultracentrifugation.

## AUTHOR CONTRIBUTIONS

Karina Kinghorn (KK) and Victoria L Bautch (VLB) conceived the project; KK, Amy Gill (AG), Feilim Mac Gabhann (FMG), and VLB designed experiments; Ferdinand le Noble (FLN) provided critical reagents and models; KK, AG, Allison Marvin (AM), Renee Li (RL), and Kaitlyn Quigley (KQ) collected data; KK and AG analyzed data; KK and VLB wrote the manuscript and all authors contributed.

## FUNDING

This work was supported by grants from the NIH-NHLBI (R35 HL139950 (VLB) and GM129074 (FMG, VLB)), a NIH-NHLBI T32 Integrated Vascular Biology Training Grant slot (5T32HL069768-18 (KK)), and an American Heart Association Predoctoral Fellowship (20PRE35080143 (KK)).

## DECLARATION OF INTERESTS

None.

## Notes

### Competing Interest Statement

The authors have declared no competing interest.

