## Supplemental Figures and Tables for "A defined clathrin-mediated trafficking pathway regulates sFLT1/VEGFR1 secretion from endothelial cells"

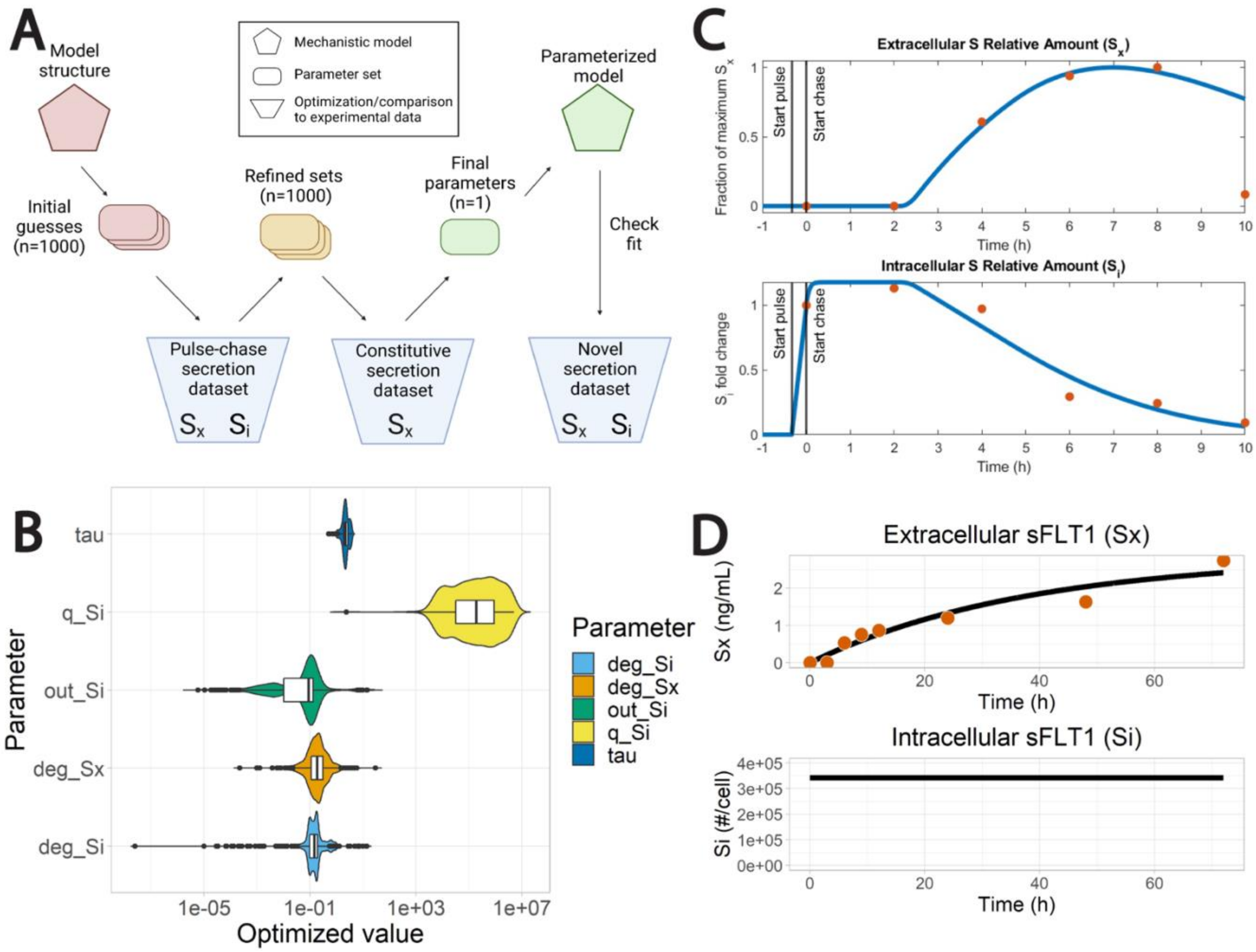


**Supplemental Fig. 1 Optimization of modeling parameters.** (A) Optimization protocol for creating a mechanistic model for sFLT1 secretion. Model parameters were first optimized to pulse-chase secretion data [55]. The lowest cost parameter set was refined to fit absolute protein concentrations from constitutive sFLT1 secretion [111]. The final model was compared to the novel sFLT1 secretion time course from this study. Schematic created with BioRender.com. (B) Distribution of values for 1000 parameter sets optimized to sFLT1 pulse-chase data [55]. All parameters except q_Si are well-constrained. (C) Experimental data (red; [55]) and simulation with best-fit optimized parameters (blue) of intracellular and extracellular sFLT1 during pulse-chase analysis of sFLT1 secretion. (D) Experimental data (red; [111]) and simulation with best-fit optimized parameters (black) of intracellular and extracellular sFLT1 during constitutive sFLT1 secretion.

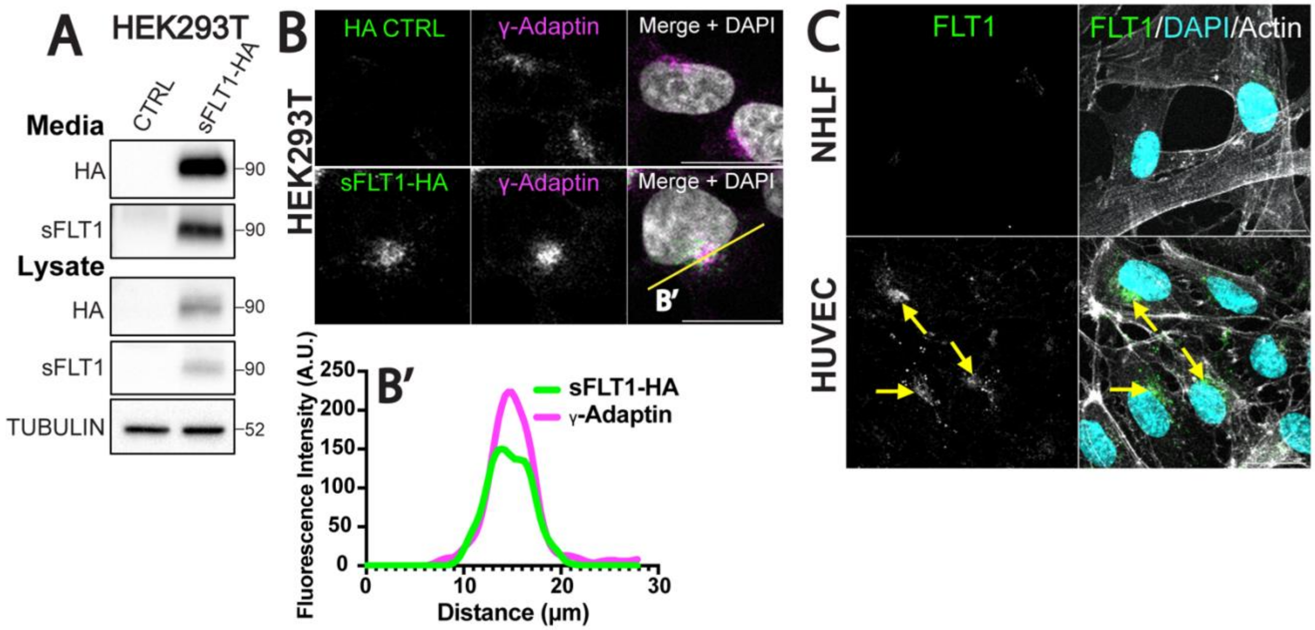


**Supplemental Fig. 2 sFLT1 and sFLT1-HA expression.** (A) Immunoblot showing sFLT1-HA in concentrated conditioned media or cell lysates of HEK293T cells transfected with indicated DNAs and probed with indicated antibodies. Tubulin loading control. (B) Immunofluorescence of HEK293T cells expressing sFLT1-HA and stained for HA, γ-Adaptin (Golgi), and DAPI (nucleus). Scale bar,20 μm. Yellow line, line scan. (B') Line scan of sFLT1-HA and γ-Adaptin fluorescence intensity. (C) Control NHLF and HUVEC immunofluorescence with indicated antibodies: FLT1, Phalloidin (actin), DAPI (nucleus). Yellow arrows: perinuclear FLT1 localization. Scale bar, 20 μm.

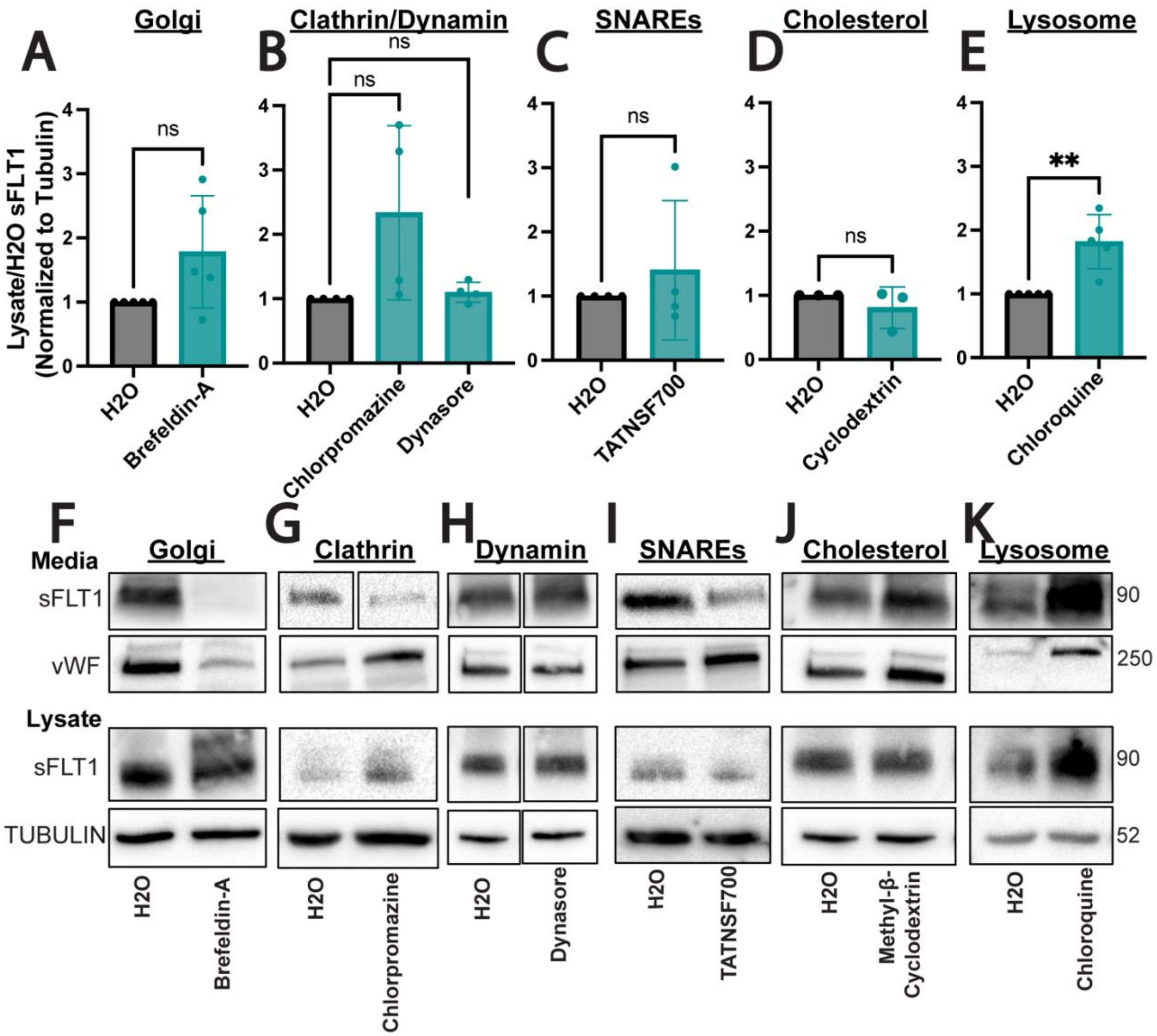


**Supplemental Fig. 3 Intracellular endothelial sFLT1 analysis and representative immunoblots after pharmacological inhibition.** (A-E) Quantification of immunoblots of HUVEC lysates treated with indicated inhibitors for 18 hr prior to collection of lysates and incubated with FLT1 antibody. Normalized to tubulin then H2O control. Mean +/- SD. Statistics: student's two-tailed t-test, **P<0.01, ns, not significant. (F-K) Representative western blots of endothelial sFLT1 and vWF in concentrated conditioned media and sFLT1 in lysates following indicated pharmacological manipulation. Tubulin loading control. Cropped bands are from the same blot.

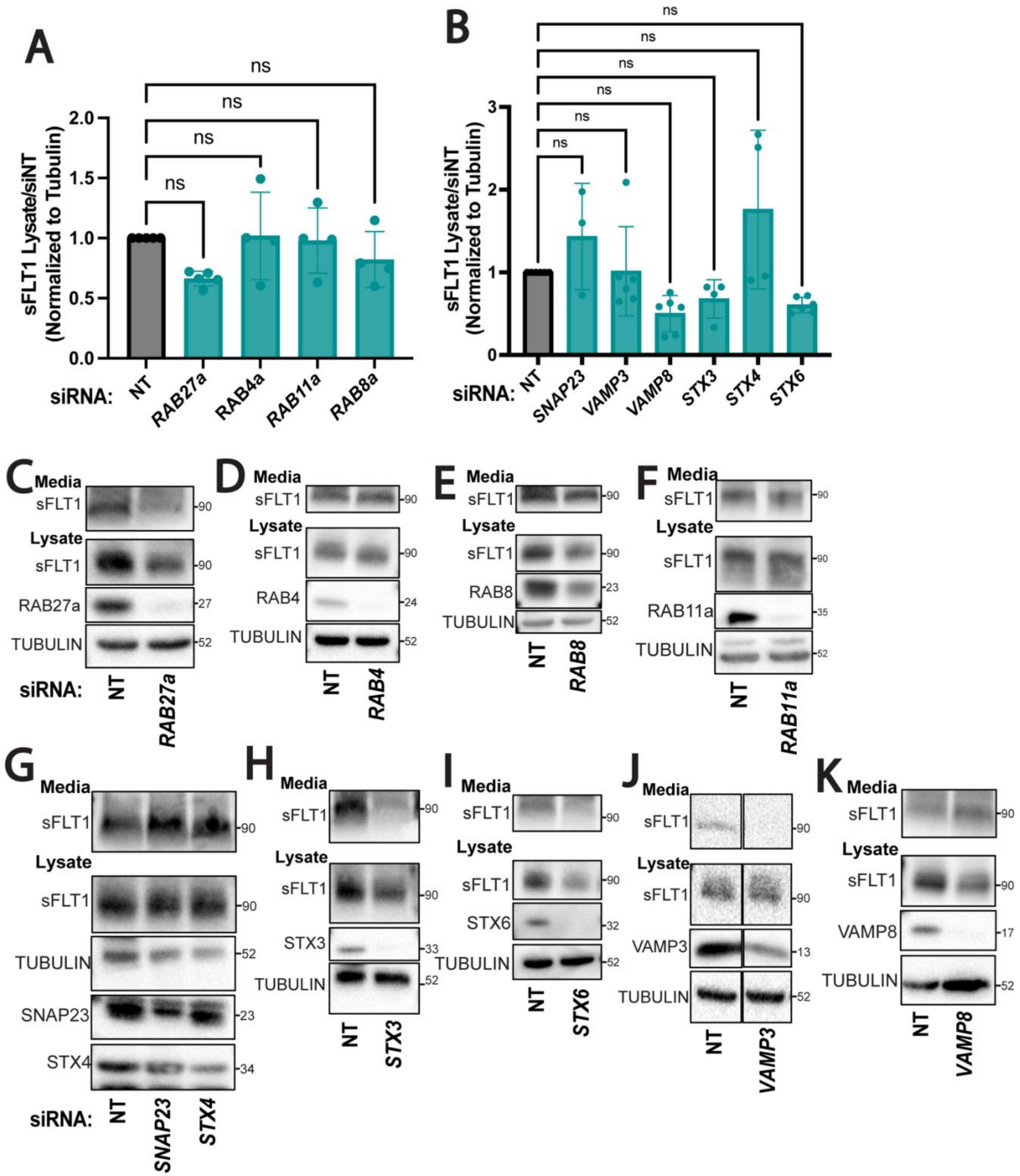


**Supplemental Fig. 4 Intracellular endothelial sFLT1 and representative western blots after RAB and SNARE manipulations.** (A,B) Quantification of immunoblots of HUVEC following indicated siRNA treatments. Normalized to tubulin then siNT. Mean +/- SD of experimental replicates shown. Statistics: student’s two-tailed t-test, ns, not significant. (C-K) Representative western blots of endothelial sFLT1 in concentrated conditioned media and lysates following indicated siRNA treatments. Tubulin loading control. Cropped bands are from the same blot.

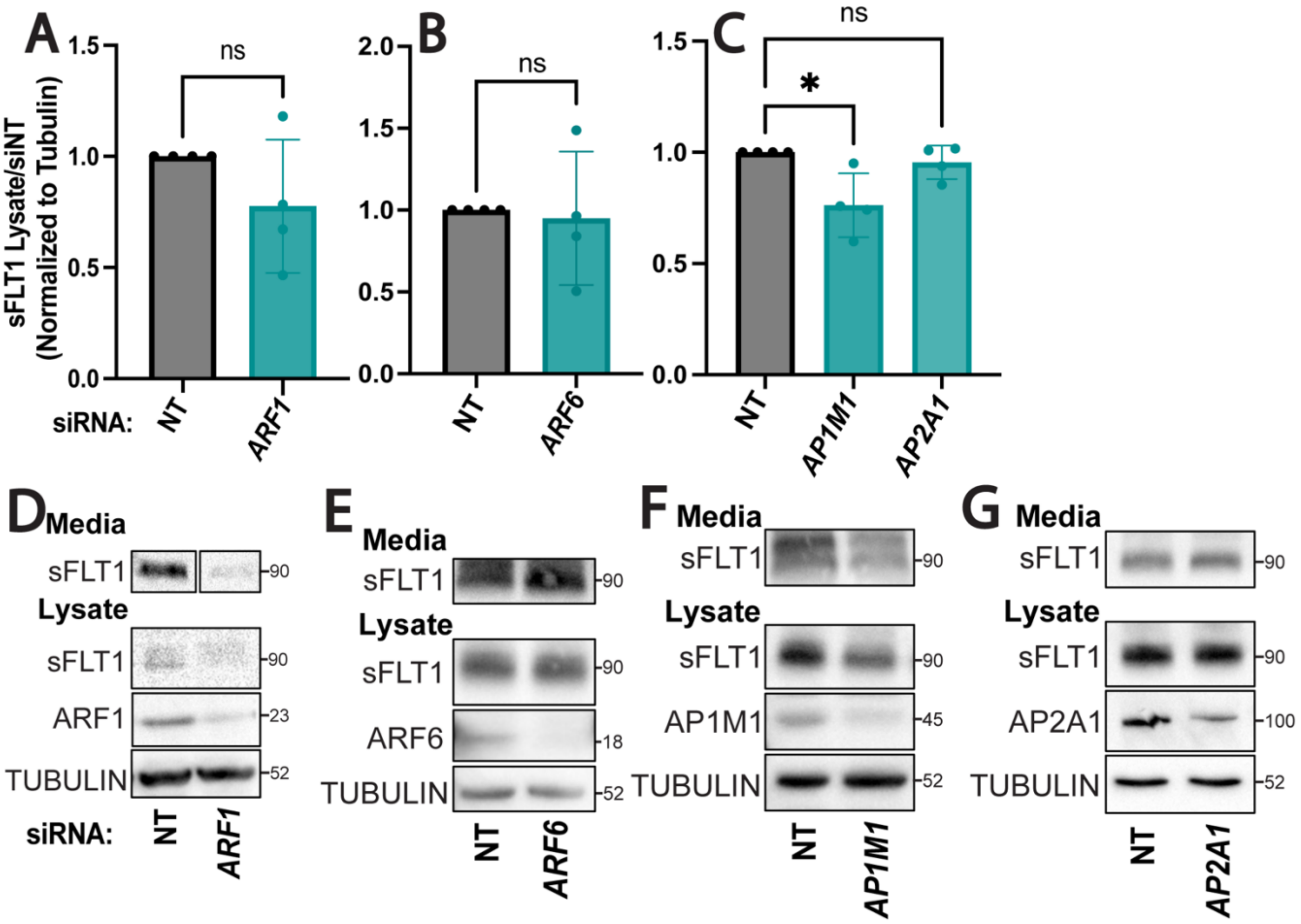


**Supplemental Fig. 5 Internal endothelial sFLT1 levels and media blots after depletion of Golgi-localized proteins.** (A-C) Quantification of immunoblots of HUVEC following indicated siRNA treatments. Normalized to tubulin then siNT. Mean +/- SD of experimental replicates shown. Statistics: student's two-tailed t-test, *P<0.05, ns, not significant. (D,G) Representative western blots of endothelial sFLT1 in concentrated conditioned media and lysates following indicated siRNA treatments. Tubulin loading control. Cropped bands are from the same blot.

### SUPPLEMENTAL TABLES

**Supplementary Table S1** Delay differential equations describing intracellular and extracellular sFLT1 amount over time in the mechanistic model. Parameter descriptions and optimal values found in Supplementary Table S4. In the $S_x$ equation, a conversion factor is applied to the $S_i$ term to obtain consistent units.

| Species | Description | Units | Rate of change equation |
|---|---|---|---|
| $S_i$ | Intracellular sFLT1 | #/cell | $\frac{dS_i(t)}{dt} = k_{q\_Si} - k_{out\_Si}S_i(t-\tau) - k_{deg\_Si}S_i(t)$ |
| $S_x$ | Extracellular sFLT1 | ng/mL | $\frac{dS_x(t)}{dt} = k_{out\_Si}S_i(t-\tau) - k_{deg\_Sx}S_x(t)$ |

**Supplementary Table S2** Characteristics of sFLT1 datasets used to build the mechanistic model.

Sx, extracellular sFLT1; Si, intracellular sFLT1; +, experimentally measured; -, not measured.

* Sx converted to fraction of maximum amount for optimization.

† Identification of a system of equations and estimation of secretion parameters.

‡ Estimation of biologically relevant production parameter values.

| Reference | Experiment | $S_x$ | $S_i$ | Measurement | Units | Time points (h) | Rationale |
|---|---|---|---|---|---|---|---|
| Jung et al., 2012 | Pulse-chase | + | + | autoradiography | Fold change* | 0, 2, 4, 6, 8, 10 | † |
| Hornig et al., 2000 | Constitutive secretion | + | - | ELISA | ng/mL | 0, 3, 6, 9, 12, 24, 48, 72 | ‡ |

**Supplementary Table S3** Input parameters for mechanistic computational model.

| Variable | Units | Value | Reference |
|---|---|---|---|
| Number of cells | cells | 5000 | Hornig et al., 2000 |
| Media volume | mL | 0.3 | Hornig et al., 2000;<br>standard media volume for 48-well plate |

**Supplementary Table S4** Optimized parameters for mechanistic computational model.

| Rate constant | Process | Units | Value |
|---|---|---|---|
| $k_{q\_Si}$ | sFLT1 synthesis | (#/cell)/h | $6.01 \times 10^4$ |
| $k_{out\_Si}$ | sFLT1 secretion | 1/h | $.0878$ |
| $k_{deg\_Si}$ | Intracellular sFLT1 degradation | 1/h | $.0873$ |
| $k_{deg\_Sx}$ | Extracellular sFLT1 degradation | 1/h | $.0264$ |
| $\tau$ | sFLT1 secretion delay | h | $2.47$ |

**Supplementary Table S5** Reported secretion rates of various proteins. † Using the following assumptions: (a) secretion rates were assumed constant when reported over a longer time span (e.g. #/cell/day) and (b) cell count was assumed constant when proliferation was not incorporated in the original rate estimate. ‡ Calculated using molecular weight and Avogadro's number. Abbreviations: ASC = adipocyte stromal cell, HUVEC = human umbilical vein endothelial cell, PBMC = peripheral blood mononuclear cell.

| Protein | Cell type | Species | Molecular weight (kDa) | Secretion rate (g/cell/h) † | Secretion rate (#/cell/h) ‡ | References |
|---|---|---|---|---|---|---|
| IgM | plasma cell | trout | 800 | $1.39 \times 10^{-10}$ | $1.05 \times 10^{8}$ | Bromage et al., 2009; Bromage et al., 2004 |
| IgE | plasma cell | human | 150 | $6.25 \times 10^{-12}$ | $2.51 \times 10^{7}$ | Werner-Favre et al., 1993 |
| IgA | plasma cell | human | 150 | $5.67 \times 10^{-12}$ | $2.27 \times 10^{7}$ | Werner-Favre et al., 1993 |
| IgG | plasma cell | human | 150 | $4.67 \times 10^{-12}$ | $1.87 \times 10^{7}$ | Werner-Favre et al., 1993 |
| IgG | plasma cell | human | 150 | $2 - 29 \times 10^{-13}$ | $0.8 - 1.2 \times 10^{7}$ | Janossy et al., 1977 |
| IgM | plasma cell | human | 950 | $1.14 \times 10^{-11}$ | $7.2 \times 10^{6}$ | Werner-Favre et al., 1993 |
| IgG | plasma cell | human | 150 | $2 - 14 \times 10^{-13}$ | $0.8 - 5.7 \times 10^{6}$ | Salmon and Smith, 1970 |
| albumin | liver | rat | 65 | $1.56 \times 10^{-13}$ | $1.44 \times 10^{6}$ | Chen and Redman, 1977 |
| IgM | plasma cell | human | 950 | $2 - 17 \times 10^{-13}$ | $0.13 - 1.0 \times 10^{6}$ | Janossy et al., 1977 |
| vWF | HUVEC | human | 250 | $2.92 \times 10^{-14}$ | $7.03 \times 10^{4}$ | Hakkert et al., 1992 |
| IFNγ | Spleen | mouse | 20 | $7.98 \times 10^{-16}$ | $4.80 \times 10^{4}$ | Favre et al., 1997 |
| TGF- β | ASC | human | 25 | $1.73 \times 10^{-17}$ | $4.17 \times 10^{3}$ | Rehman et al., 2004 |
| IL-6 | PBMC | human | 21 | $7.8 - 10 \times 10^{-17}$ | $2.2 - 2.9 \times 10^{3}$ | O'Mahony et al., 1998 |
| TNF-α | PBMC | human | 17 | $3.5 - 5.4 \times 10^{-17}$ | $1.2 - 1.9 \times 10^{3}$ | O'Mahony et al., 1998 |
| HGF | ASC | human | 80 | $1.71 \times 10^{-16}$ | $1.28 \times 10^{3}$ | Rehman et al., 2004 |
| VEGF (hypoxia) | ASC | human | 40 | $8.31 \times 10^{-17}$ | $1.25 \times 10^{3}$ | Rehman et al., 2004 |
| IL-1β | PBMC | human | 17 | $2.1 - 2.2 \times 10^{-17}$ | $7.4 - 7.9 \times 10^{2}$ | O'Mahony et al., 1998 |
| VEGF (normoxia) | ASC | human | 40 | $1.67 \times 10^{-17}$ | $2.51 \times 10^{2}$ | Rehman et al., 2004 |
| bFGF | ASC | human | 22 | $1.72 \times 10^{-17}$ | $4.71 \times 10^{1}$ | Rehman et al., 2004 |
| GM-CSF | ASC | human | 15 | $1.17 \times 10^{-17}$ | $4.68 \times 10^{1}$ | Rehman et al., 2004 |

**Key Resources**

**Supplemental Table S6** Pharmacological Inhibitors.

| Inhibitor | Final Concentration | Stock Solution | Company | Target |
|---|---|---|---|---|
| Brefeldin-A | 1 mg/mL | 5 mg/mL in DMSO | Biolegend (420601) | Blocks the GEF of ARF1 and leads to Golgi collapse |
| Chlorpromazine | 10 μM (HUVEC), 100 μM (fish) | 10 mM in H2O | Sigma (C8138) | Prevents assembly and disassembly of clathrin lattices |
| Dynasore | 5 μM | 10 mM in DMSO | Santa Cruz (sc-202592) | Inhibits the GTPase of dynamin |
| TATNSF-700 | 1 μM | 1 mM in H2O | AnaSpec (AS-62238) | Competes for the NSF binding and blocks SNARE disassembly |
| Methyl-B-Cyclodextrin | 1mM | 10 mM in H2O | Sigma (C4555) | Depletes membranes of cholesterol |
| Chloroquine | 10 μg/mL | 10 mg/mL in H2O | Sigma (C6628) | Elevates lysosomal pH to block degradation |

**Supplemental Table S7.** siRNAs.

| siRNAs | Catalog # | Company | pmol/10cm2 | Target Sequence (5'--> 3') |
|---|---|---|---|---|
| FLT1-1 | 4392420 (s5287) | ThermoFisher | 200 pmol | GGUGAGUAAGGAAAGCGAAtt |
| FLT1-2 | sc-29319 | Santa Cruz | 200 pmol | N/A |
| sFLT1 | 4390827 (custom) | ThermoFisher | 200 pmol | AAGGCUGUUUUCUCUCGGAUU |
| mFLT1 | 4390827 (custom) | ThermoFisher | 200 pmol | AAAUAGUGGGUUUACAUACUU |
| ARF1-1 | 4390824 (s1551) | ThermoFisher | 200 pmol | CCAUAGGCUUCAACGUGGAtt |
| ARF1-2 | L-011580-00-0005 | Dharmacon | 200 pmol | UGACAGAGAGCGUGUGAAC, CGGCCGAGAUCACAGACAA, ACGAUCCUCUACAAGCUUA, GAACCAGAAGUGAACGCGA |
| STX6-1 | 4392420 (s19959) | ThermoFisher | 200 pmol | GCAACUGAAUUGAGUAUAAtt |

| | | | | |
|---|---|---|---|---|
| STX6-2 | 4392420 (s19958) | ThermoFisher | 200 pmol | CCAACGAGCUGAGAAAUAAtt |
| AP1M1 | 4392420 (s17032) | ThermoFisher | 200 pmol | ACAACUUUGUUAUCAUCUAtt |
| AP2A1 | 4390824 (s184) | ThermoFisher | 200 pmol | GCCGAUGAGUUGCUGAAUAtt |
| RAB27a-1 | 4390824 (s11695) | ThermoFisher | 200 pmol | GGAAGACCAGUGUACUUUAtt |
| RAB27a-2 | 4390824 (s11693) | ThermoFisher | 200 pmol | GCCUCUACGGAUCAGUUAAtt |
| RAB4 | 439084 (s11675) | ThermoFisher | 200 pmol | GGUCCGUGACGAGAAGUUAtt |
| RAB11a-1 | sc-36340 | Santa Cruz | 200 pmol | N/A |
| RAB11a-2 | 4390824 (s16702) | ThermoFisher | 200 pmol | CAACAAUGUGGUUCCUAUUtt |
| RAB8a | 4390824 (s8681) | ThermoFisher | 200 pmol | CUUUAAAAUUAGGACCAUAtt |
| ARF6-1 | 4390824 (s1567) | ThermoFisher | 200 pmol | CCAAGGUCUCAUCUUCGUAtt |
| ARF6-2 | 4390824 (s1565) | ThermoFisher | 200 pmol | CUCUCAUCUUCGUAGUGGAtt |
| SNAP23-1 | sc-41308 | Santa Cruz | 200 pmol | N/A |
| SNAP23-2 | 4392420 (s16709) | ThermoFisher | 200 pmol | GGAACAACUAAACCGCAUAtt |
| STX4-1 | sc-36590 | Santa Cruz | 200 pmol | N/A |
| STX4-2 | 4392420 (s13597) | ThermoFisher | 200 pmol | UGAUCAAUCGGAUUGAGAAtt |
| STX3-1 | sc-616132 | Santa Cruz | 200 pmol | N/A |
| STX3-2 | 4392420 (s13592) | ThermoFisher | 200 pmol | GGCACGAGAUGAAACGAAAtt |
| VAMP3-1 | sc-41338 | Santa Cruz | 200 pmol | N/A |
| VAMP3-2 | 4392420 (s17856) | ThermoFisher | 200 pmol | CGGGAUUACUGUUCUGGUUtt |
| VAMP8-1 | sc-41300 | Santa Cruz | 200 pmol | N/A |
| VAMP8-2 | 4392420 (s16522) | ThermoFisher | 200 pmol | GGAGUUAAGAAUAUUAUGAtt |
| Non-targeting-1 | 4390844 | ThermoFisher | 200 pmol | N/A |
| Non-targeting-2 | 4390847 | ThermoFisher | 200 pmol | N/A |
| Non-targeting-3 | sc-37007 | Santa Cruz | 200 pmol | N/A |

**Supplemental Table S8** Primary Antibodies.

| Target | Host Species | Company | Catalog Number | Western Dilution | IF Dilution |
|---|---|---|---|---|---|
| FLT1 | goat | RnD | AF321 | 1:10,000 | 1:500 |
| VWF | mouse | ThermoFisher | MA5-14029 | 1:500 | 1:1000 |
| HA | rabbit | Cell Signaling | 3724S | 1:1,000 | 1:500 |
| HA | mouse | Biolegend | 901502 | 1:500 | 1:500 |
| GM130 (Cis-Golgi) | rabbit | abcam | ab52649 | 1:1,2000 | 1:500 |
| Gamma-Adaptin (Trans-Golgi) | rabbit | abcam | ab220251 | N/A | 1:500 |
| STX6 (Syntaxin 6, Trans-Golgi) | rabbit | Cell Signlaing | 2869s | 1:500 | 1:250 |
| AP1M1 (adaptor protein 1) | rabbit | Proteintech | 12112-1-AP | 1:1000 | N/A |
| AP2A1 | mouse | ThermoFisher | MA3-061 | 1:2000 | N/A |
| Clathrin Heavy Chain | mouse | ThermoFisher | MA1-065 | N/A | 1:500 |
| Tubulin | rabbit | Cell Signaling | 2144S | 1:50,000 | N/A |
| Tubulin | mouse | Cell Signaling | 3873S | 1:100,000 | N/A |
| RAB27a | rabbit | Cell Signaling | 95394S | 1:2000 | 1:250 |
| ARF1 | rabbit | abcam | ab76082 | 1:1000 | N/A |
| ARF6 | rabbit | abcam | ab77581 | 1:1000 | N/A |
| SNAP23 | rabbit | abcam | ab3340 | 1:2000 | N/A |
| STX3 | rabbit | abcam | ab133750 | 1:2000 | N/A |
| STX4 | rabbit | abcam | ab184545 | 1:1000 | N/A |
| VAMP3 (Cellubrevin) | rabbit | abcam | ab43080 | 1:5000 | N/A |
| VAMP8 | rabbit | abcam | ab76021 | 1:10,000 | N/A |
| RAB4 | mouse | ThermoFisher | MA5-17161 | 1:1000 | N/A |
| RAB11 | rabbit | abcam | ab65200 | 1:1000 | N/A |

**Supplemental Table S9** Secondary Antibodies.

| Antibody | Catalog Number | Company | Western Dilution | IF Dilution |
|---|---|---|---|---|
| Donkey anti-goat IgG HRP | PA1-28664 | ThermoFisher | 1:20,000 | N/A |
| Donkey anti-Rabbit IgG (H+L) Highly Cross-Adsorbed Secondary Antibody, HRP | A16035 | ThermoFisher | 1:10,000 | N/A |
| Donkey anti-Mouse IgG (H+L) Highly Cross-Adsorbed Secondary Antibody, HRP | A16011 | ThermoFisher | 1:10,000 | N/A |
| Goat anti-Rabbit HRP | 31460 | ThermoFisher | 1:10,000 | N/A |
| Goat anti-Mouse HRP | 31430 | ThermoFisher | 1:10,000 | N/A |
| Donkey anti Goat IgG (H+L) Highly Cross Absorbed Secondary Antibody, Alexa Fluor 488 | A-11055 | ThermoFisher | N/A | 1:1,000 |
| Donkey anti Rabbit IgG (H+L) Highly Cross Absorbed Secondary Antibody, Alexa Fluor 488 | A-21206 | ThermoFisher | N/A | 1:1,000 |
| Donkey anti Mouse IgG (H+L) Highly Cross Absorbed Secondary Antibody, Alexa Fluor 488 | A-21202 | ThermoFisher | N/A | 1:1,000 |
| Donkey anti Rabbit IgG (H+L) Highly Cross Absorbed Secondary Antibody, Alexa Fluor 594 | A-21207 | ThermoFisher | N/A | 1:1,000 |
| Donkey anti Mouse IgG (H+L) Highly Cross Absorbed Secondary Antibody, Alexa Fluor 594 | A-21203 | ThermoFisher | N/A | 1:1,000 |
| Donkey anti Goat IgG (H+L) Highly Cross Absorbed Secondary Antibody, Alexa Fluor 647 | A32849 | ThermoFisher | N/A | 1:1,000 |
| Donkey anti Rabbit IgG (H+L) Highly Cross Absorbed Secondary Antibody, Alexa Fluor 647 | A-31573 | ThermoFisher | N/A | 1:1,000 |
| Donkey anti Mouse IgG (H+L) Highly Cross Absorbed Secondary Antibody, Alexa Fluor 647 | A-31571 | ThermoFisher | N/A | 1:1,000 |
| Alexa Fluor 594 Phalloidin | A12381 | ThermoFisher | N/A | 1:500 |
| Alexa Fluor 647 Phalloidin | A22287 | ThermoFisher | N/A | 1:500 |
| DAPI | 10236276001 | Sigma | N/A | 1:1,000 |
